# A widespread family of viral sponge proteins reveals specific inhibition of nucleotide signals in anti-phage defense

**DOI:** 10.1101/2024.12.30.630793

**Authors:** Renee B. Chang, Hunter C. Toyoda, Samuel J. Hobbs, Desmond Richmond-Buccola, Tanita Wein, Nils Burger, Edward T. Chouchani, Rotem Sorek, Philip J. Kranzusch

## Abstract

Cyclic oligonucleotide-based antiviral signaling systems (CBASS) are bacterial anti-phage defense operons that use nucleotide signals to control immune activation. Here we biochemically screen 57 diverse *E. coli* and *Bacillus* phages for the ability to disrupt CBASS immunity and discover anti-CBASS 4 (Acb4) from the *Bacillus* phage SPO1 as the founding member of a large family of >1,300 immune evasion proteins. A 2.1 Å crystal structure of Acb4 in complex with 3′3′-cGAMP reveals a tetrameric assembly that functions as a sponge to sequester CBASS signals and inhibit immune activation. We demonstrate Acb4 alone is sufficient to disrupt CBASS activation *in vitro* and enable immune evasion *in vivo*. Analyzing phages that infect diverse bacteria, we explain how Acb4 selectively targets nucleotide signals in host defense and avoids disruption of cellular homeostasis. Together, our results reveal principles of immune evasion protein evolution and explain a major mechanism phages use to inhibit host immunity.

## Introduction

The ability to sense and respond to viral infection is an essential cellular process conserved across domains of life. In bacteria, a major mechanism of antiviral immunity is synthesis of nucleotide immune signals that activate defense against bacteriophage (phage) infection.^1,2^ Common bacterial nucleotide immune signaling pathways include cyclic oligonucleotide-based anti-phage signaling system (CBASS), Thoeris, Pycsar, and Type III CRISPR operons.^3–9^ The most abundant of these systems is CBASS, occurring in >16% of all sequenced bacteria.^10–13^ CBASS operons encode two main components: a cGAS/DncV-like nucleotidyltransferase (CD-NTase) enzyme that responds to phage infection and catalyzes the synthesis of a nucleotide immune signal, and a CD-NTase-associated protein (Cap) effector that recognizes the nucleotide immune signal and induces cell death or cell stasis to prevent phage propagation.^3,4,13–17^ CD-NTase enzymes can synthesize distinct nucleotide immune signals including cyclic di- or trinucleotide species, signals composed of all four common purine and pyrimidine bases, and signals containing 3′–5′ and 2′–5′ phosphodiester linkages.^2,4,14^ CBASS operons additionally encode an enormous diversity of Cap effectors including nucleases, NAD^+^-degrading enzymes, and membrane-disrupting proteins that create a strong barrier to phage replication.^3,4,10,14,16,18–20^

The arms race between viruses and hosts has led to evolution of distinct viral immune evasion strategies that target host nucleotide immune signaling. First, phages encode dedicated nuclease enzymes that rapidly degrade nucleotide immune signals including anti-CBASS 1 (Acb1) that inhibits CBASS, anti-Pycsar 1 (Apyc1) that inhibits Pycsar, and anti-CRISPR III-1 (AcrIII-1) that inhibits Type III CRISPR immunity.^21,22^ Nucleotide immune signal degrading enzymes are also encoded in animal viruses including eukaryotic viral homologs of Acb1 and a family of nucleases named poxins that inhibit mammalian and insect innate immunity.^23–26^ Second, phages encode sponge proteins that tightly bind and sequester nucleotide immune signals to inhibit downstream effector activation.^27–30^ Three major families of viral sponge immune evasion proteins are defined by anti-CBASS 2 (Acb2), Thoeris anti-defense 1 (Tad1), and Thoeris anti-defense 2 (Tad2). Recent results demonstrate that individual sponge proteins can evolve to target a broad range of signals in CBASS and Thoeris anti-phage defense, highlighting the importance of suppressing host nucleotide immune signaling to ensure successful phage replication.^27–33^

The emergence of sponge proteins as a conserved strategy of viral immune evasion creates several open questions. Currently, sponge protein identification has been limited to genetic based methods screening for phages with altered susceptibility to host anti-phage defense.^27–30^ Each family of sponge protein is structurally distinct, suggesting that an additional diversity of sponge proteins remain to be discovered. Structural studies of viral sponges have identified the ligand-binding pockets that enable nucleotide immune signal sequestration^27–33^, but it remains unclear how sponges adapt ligand specificity to efficiently sequester nucleotide immune signals while avoiding non-relevant signals in the host environment. Finally, the evolutionary origin of sponge proteins and related mechanisms of viral immune evasion are unknown, limiting understanding of how viruses acquire mechanisms to inhibit host antiviral defense.

Here we develop a forward biochemical approach for the discovery of viral sponges from infected lysates. We apply our unbiased method to screen a diverse collection of 57 phages and identify *Bacillus* phage SPO1 Acb4 as the founding member of a family of >1,300 anti-CBASS sponges. We show that Acb4 expression is sufficient to overcome CBASS defense *in vivo*, underscoring its role as an immune evasion protein. Our structural analyses of Acb4 reveal a tetrameric assembly that sequesters CBASS immune signals through nucleobase-specific pockets, providing insight into how viral sponge proteins enable specific recognition of immune signals. In contrast to Acb1 and Acb2, which are relatively rare phage proteins restricted to specific groups of phages^21,28^, our analyses reveal Acb4 as the largest known family of anti-CBASS proteins to date and provides insight into how viruses can overcome prevalent CBASS systems encoded across diverse bacterial phyla. Finally, phylogenetic and biochemical analyses of Acb4 homologs demonstrate that viral sponges can arise from common bacterial host protein domains and explain how further adaptation enables sponges to avoid recognizing signaling molecules required for cell homeostasis. Together, our results reveal Acb4 as a widespread family of viral sponge proteins and provide a framework for understanding how phages employ viral sponges to selectively inhibit nucleotide immune signals to evade host defense.

## Results

### Discovery of SPO1 Acb4 as a phage-encoded sponge

To discover viral proteins that inhibit bacterial anti-phage defense, we developed a biochemical screen capable of detecting viral nucleotide immune signal sponges directly from phage-infected lysates. Using radiolabeled 3′3′-cyclic GMP-AMP (3′3′-cGAMP), a common second messenger signal in CBASS immunity^3,4^, we optimized an electrophoretic mobility shift assay (EMSA) to measure the ability of phage proteins in infected cell lysates to sequester or degrade host nucleotide immune signals (Figures 1A and S1A). To validate our approach, we tested lysates infected with the *E. coli* phage T4 which encodes two distinct anti-CBASS proteins, Acb1 and Acb2 (Figure S1A).^21,28,29^ Acb1 is a nuclease that disrupts CBASS-mediated immunity by degrading 3′3′-cGAMP and related nucleotide immune signals, and Acb2 is a sponge that sequesters 3′3′-cGAMP to prevent downstream effector activation. In the presence of cell lysates infected with wildtype phage T4, we observed that phage Acb1 activity is dominant and results in complete degradation of radiolabeled 3′3′-cGAMP that appears as faster-moving species on the gel (Figure S1A). Using a mutant phage T4 virus lacking Acb1 (T4 Δ*acb1*)^21^, we observed that 3′3′-cGAMP remains intact and is shifted upwards as a slower-moving species in the gel due to interaction with the sponge protein Acb2 (Figure S1A). We confirmed that all 3′3′-cGAMP degradation and binding activity is lost in the presence of cell lysates infected with a phage T4 virus lacking both Acb1 and Acb2 (T4 Δ*acb1* Δ*acb2*)^26^ (Figure S1A). These results demonstrate our biochemical assay allows for robust detection of phage anti-CBASS immune evasion proteins that bind or degrade host nucleotide immune signals.

**Figure 1.**
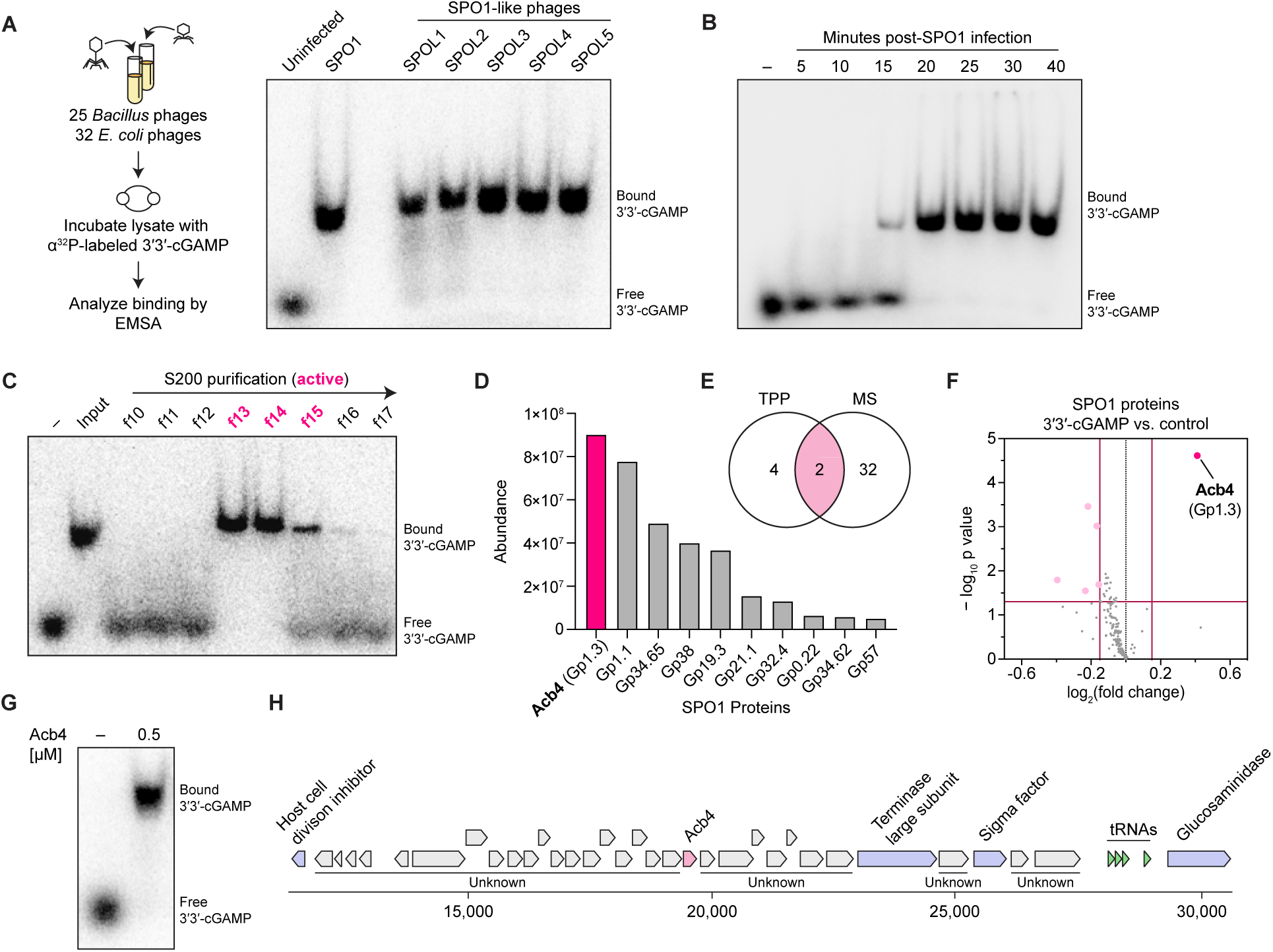
Discovery of SPO1 Acb4 as a 3′3′-cGAMP binding protein. (A) Schematic of biochemical screen for discovery of 3′3′-cGAMP binding activity in phage-infected lysates using electrophoretic mobility shift assay (EMSA). Representative EMSA depicting 3′3′-cGAMP binding activity following infection with SPO1 or SPO1-like phages (SPOL1–SPOL5). Data are representative of at least *n* = 3 independent experiments. Phages used in the screen are listed in Table S1, and primary data are shown in Figure S1C. (B) EMSA analysis depicting the time course of 3′3′-cGAMP binding activity in SPO1-infected lysates. SPO1 infection experiments were performed at 37°C and data are representative of *n* = 2 independent experiments. (C) Representative EMSA analysis demonstrating enrichment of candidate Acb4 genes by activity-guided fractionation. SPO1-infected lysates were fractionated using hydrophobic interaction chromatography followed by S200 size-exclusion chromatography. Fractions enriched in 3′3′-cGAMP binding activity were analyzed by mass spectrometry (MS) to identify candidate SPO1 proteins. For additional details on purification scheme, see Figure S2A. (D) MS analysis of SPO1 proteins detected in a fraction with peak 3′3′-cGAMP binding activity following biochemical fractionation. Abundance was determined by normalizing the sum intensity of each SPO1 protein to the total number of detected peptides. Complete list of SPO1 proteins identified in the selected active fraction are shown in Figure S2B. (E) Comparison of candidate Acb4 proteins identified independently by thermal proteome profiling (TPP), or biochemical fractionation followed by MS. For a detailed list of proteins identified by each approach, see Figure S2B. (F) Volcano plot depicting SPO1 proteins that undergo a shift in thermal stability upon treatment with 100 µM 3′3′-cGAMP. Data are representative of *n =* 5 independent replicates for control and 3′3′-cGAMP-treated samples. (G) EMSA analysis of 3′3′-cGAMP binding with recombinant Acb4 produced in *Escherichia coli.* Data are representative of at least *n* = 3 independent experiments. For details on recombinant Acb4 expression and purification, refer to Figures S2C and S2D. (H) Schematic illustrating genes neighboring SPO1 *acb4*.

We next applied this approach to screen a diverse collection of 57 phages that infect *E. coli* or *B. subtilis* cells (Figures 1A, S1C, and Table S1). As a positive hit from our screen, we observed that cell lysates infected with the *B. subtilis* phage SPO1 completely shifted 3′3′-cGAMP (Figure 1A). Phage SPO1 encodes an anti-Thoeris sponge named Tad2 that sequesters the Thoeris signaling molecule 1′′–3′ gcADPR.^30,32^ We purified recombinant SPO1 Tad2 and observed that this protein does not interact with 3′3′-cGAMP and is unable to explain the altered migration of 3′3′-cGAMP in phage SPO1 infected cell lysates (Figure S1B). We tested five related SPO1-like phages (SPOL1–SPOL5) from the subfamily *Spounavirinae* and observed that each infected cell lysate retained the ability to shift 3′3′-cGAMP (Figure 1A). Similar to the expression pattern of phage T4 Acb1^21^, phage SPO1 3′3′-cGAMP-binding activity occurs early at ∼15 minutes post-infection and is maintained throughout the later stages of viral replication, suggesting that *Spounavirinae* phages encode a conserved nucleotide immune signal sponge that inhibits CBASS anti-phage defense (Figure 1B).

To determine the phage SPO1 protein responsible for 3′3′-cGAMP-binding, we next used an activity-guided fractionation and mass spectrometry approach to identify candidate Acb proteins from infected cell lysates (Figures 1C–E, S2A, and S2B). In parallel, we performed thermal proteome profiling^34–36^ to define phage SPO1 proteins that are specifically stabilized in the presence of 3′3′-cGAMP (Figure 1F). Strikingly, each approach independently revealed the same top viral protein hit as the uncharacterized phage SPO1 protein gp1.3 (Figures 1 D–F). Recombinant expression and purification of phage SPO1 gp1.3 confirmed 3′3′-cGAMP binding activity, and we named this protein anti-CBASS 4 (Acb4) (Figures 1G, 1H, and S2C–E). The *acb4* gene is encoded in the phage SPO1 and phage SPOL1–L5 genomes within a cluster of small uncharacterized proteins (Figure 1H), in agreement with the known tendency of anti-CBASS genes to reside in putative phage anti-defense islands.^21,28,37,38^ Together, these results reveal Acb4 as a nucleotide immune signal sponge conserved in SPO1 and SPO1-like phages.

### Acb4 subverts host CBASS immunity

We next measured the ability of Acb4 to inhibit CBASS signaling *in vitro* and *in vivo* during phage infection. CD-NTase-associated protein 5 (Cap5) is a prevalent CBASS nuclease effector protein that is activated by nucleotide immune signaling and induces host cell death through non-specific dsDNA degradation.^14^ We reconstituted *Burkholderia pseudomallei* Cap5 activation and dsDNA degradation *in vitro* using purified components and observed that Acb4 is sufficient to strongly suppress 3′3′-cGAMP signaling and CBASS effector activation (Figure 2A). The genomes of SPO1-like phages contain heavily modified DNA bases that limit methods for genetic manipulation.^39,40^ We therefore tested the function of SPO1 Acb4 *in vivo* using a panel of phage T4 chimeric variants. Beginning with a phage T4 variant with a deletion in *acb1*^21^, we engineered a chimeric virus where the phage T4 *acb2* gene is replaced with phage SPO1 *acb4* and confirmed generation of the recombinant phage T4 Δ*acb1/* Δ*acb2::acb4* virus by sequencing (Figure 2B and S3A). *E. coli* lysates from phage T4 Δ*acb1/* Δ*acb2::acb4* infected cells regain the ability to bind radiolabeled 3′3′-cGAMP, demonstrating that phage SPO1 Acb4 protein is actively expressed and functional in the setting of phage infection (Figure S3B).

**Figure 2.**
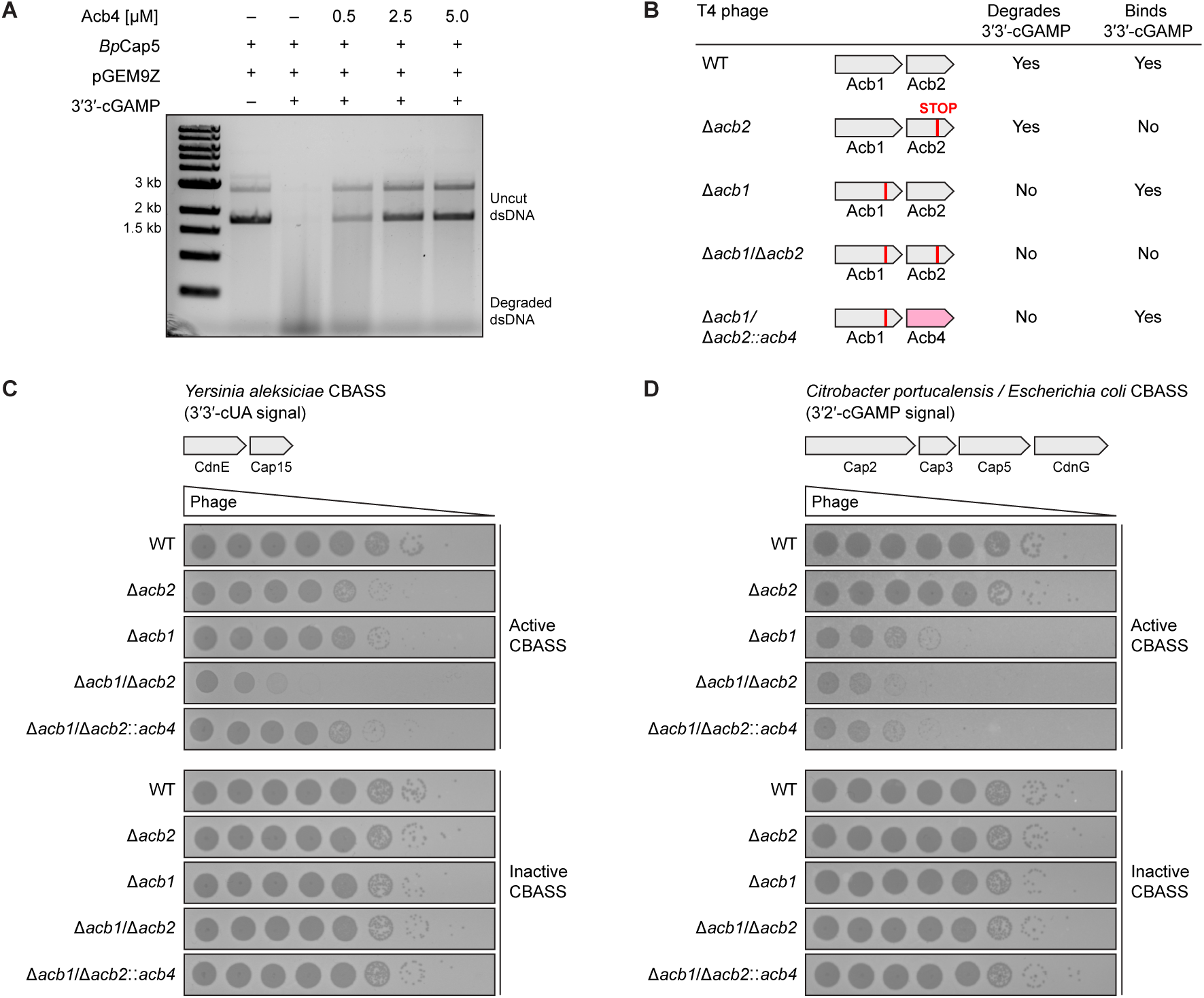
Acb4 subverts host CBASS immunity. (A) DNA cleavage analysis of uncut plasmid DNA (pGEM9Z) incubated with purified *Burkholderia pseudomallei* Cap5 and 3′3′-cGAMP pre-treated with a titration (500 nM–5 μM) of recombinant Acb4. Data are representative of at least *n* = 3 independent experiments. (B) Summary of 3′3′-cGAMP binding and degradation properties in wildtype phage T4 and engineered phage T4 variants. (C) Representative plaque assay performed using *Escherichia coli* BL21 cells harboring active or inactive forms of a CBASS operon from *Yersinia aleksiciae* and challenged with T4 phages in Figure 2B. Data are representative of *n* = 3 independent experiments. (D) Representative plaque assay performed using *Escherichia coli* BL21 cells harboring active or inactive forms of a CBASS operon from *Citrobacter portucalensis* / *Escherichia coli* and challenged with T4 phages in Figure 2B. Data are representative of *n* = 3 independent experiments.

We next compared replication fitness of the panel of the recombinant phage variants in *E. coli* cells expressing well-characterized CBASS operons from *Yersinia aleksiciae* (a Type I CdnE operon that signals through 3′3′-cUA) and *Citrobacter portucalensis / Escherichia coli* UCI 32 (a Type II CdnG operon that signals through 3′2′-cGAMP) that are known to inhibit phage T4 replication.^14,19,41^ We confirmed *in vitro* that phage SPO1 Acb4 is capable of sequestering both 3′3′-cUA and 3′2′-cGAMP CBASS signals (discussed below). In the presence of active CBASS defense, the replication of phage T4 lacking Acb1 and Acb2 (T4 Δ*acb1/*Δ*acb2*) was severely compromised with ∼10,000× fold loss in viral fitness (Figures 2C and 2D). Strikingly, phage T4 variants expressing SPO1 Acb4 as the only anti-CBASS protein (T4 Δ*acb1/*Δ*acb2::acb4*) subverted *Yersinia aleksiciae* CBASS defense and re-gained robust viral replication (Figure 2C). The T4 Δ*acb1/*Δ*acb2::acb4* and T4 Δ*acb1* viruses grew to equal titers demonstrating that either viral sponge protein Acb4 or Acb2 is alone sufficient to inhibit *Yersinia aleksiciae* CBASS defense (Figure 2C). Interestingly, successful phage T4 replication in the presence of *Citrobacter portucalensis* / *Escherichia coli* CBASS immunity was predominantly dependent on the presence of the Acb1 nuclease protein (Figure 2D). Neither Acb4 nor Acb2 sponge proteins were alone sufficient to inhibit *Citrobacter portucalensis / Escherichia coli* CBASS defense, suggesting that in some contexts nuclease and sponge anti-CBASS proteins may have distinct advantages in mediating viral immune evasion and explaining why viruses like phage T4 have evolved multiple distinct immune evasion strategies in parallel (Figure 2D). Together, these results demonstrate that Acb4 is an anti-CBASS protein sufficient to rescue viral replication and subvert host anti-phage defense.

### Structural basis of Acb4 assembly and nucleotide immune signal recognition

To define the molecular mechanism of Acb4 nucleotide sequestration and CBASS evasion, we determined a 2.10 Å crystal structure of the phage SPO1 Acb4–3′3′-cGAMP complex (Figure 3A and Table S2, Crystallographic Statistics). Acb4 is a small 96 amino acid protein that assembles into a tetrameric pinwheel-like structure (Figures 3A, 3B, and S2E). Each Acb4 protomer is composed of a core region of three antiparallel β-strands (β_1_–β_3_) flanked by an N-terminal helical region (α_1_–α_3_) and a longer C-terminal extension with two helices (α_4_–α_5_) (Figure S4A). Individual Acb4 protomers form two distinct protein–protein interfaces that interlock to create the tetrameric pinwheel-like assembly. First, the β_1_–β_3_ central region of each Acb4 protomer aligns head-to-tail against the central region of a partnering subunit in a beta-sandwich interaction that buries ∼143 Å^2^ of hydrophobic surface area and creates a dimeric unit (Figure 3B). Second, dimeric units clamp together at a “joint” interface formed between the C-terminal end of helix α_4_ from each Acb4 protomer and a cleft between helix α_4_ and the β_2_–β_3_ loop of Acb4 protomers in the opposite dimeric unit (Figure 3B). Each Acb4 protein-protein interface is highly conserved in phage SPO1 and SPO1-like Acb4 proteins, and we confirmed using SEC-MALS analysis that *apo* Acb4 forms a stable tetrameric assembly in solution (Figure S2E). Notably, Acb4 exhibits no structural homology to any previously characterized sponge immune evasion protein (i.e., Acb2, Tad1, and Tad2) or nucleotide binding protein (Figure S5).

**Figure 3.**
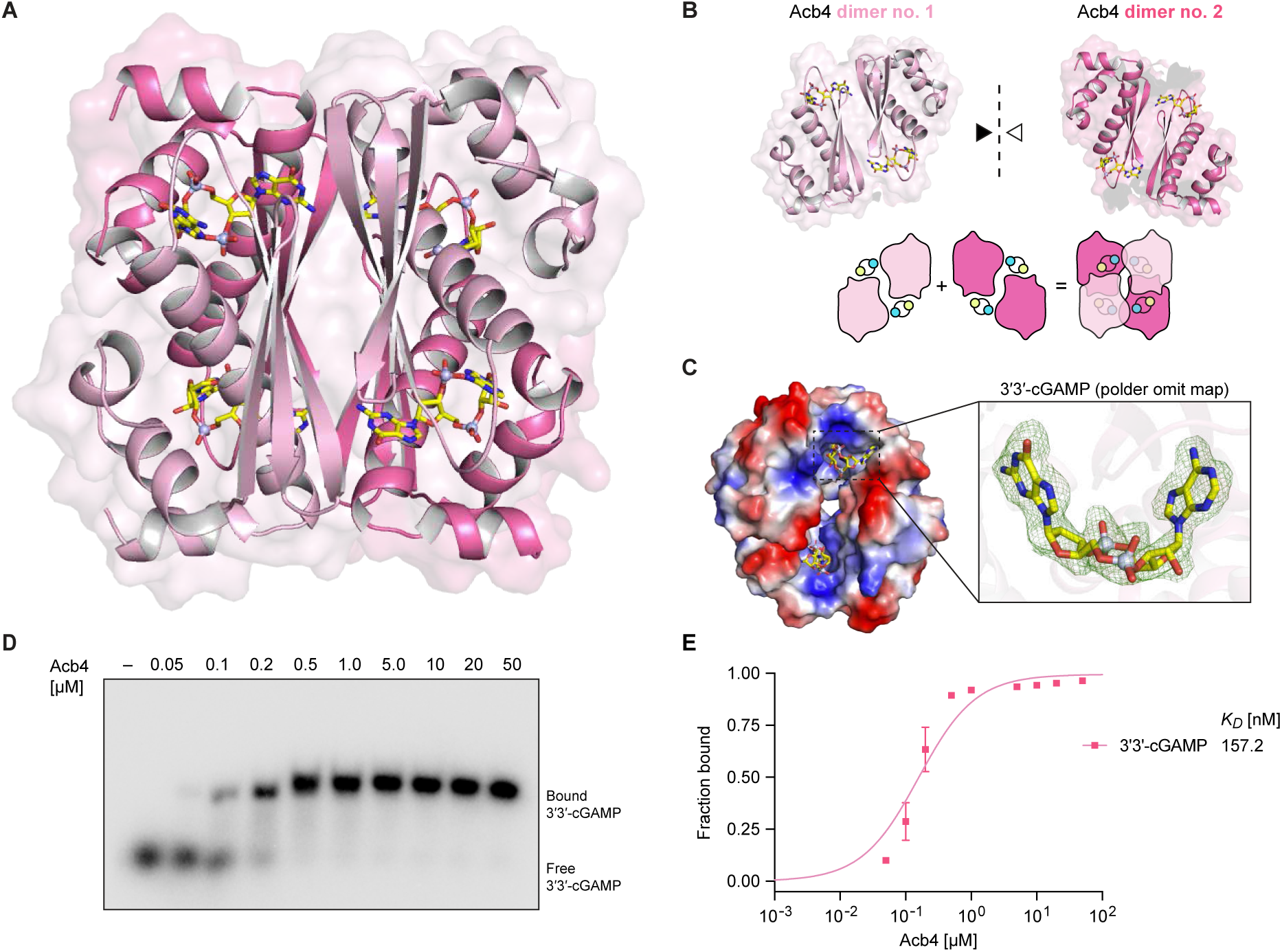
Acb4 forms a tetrameric assembly that sequesters four molecules of 3′3′-cGAMP. (A) Co-crystal structure of Acb4 from *Bacillus* phage SPO1 in complex with 3′3′-cGAMP. (B) Views of Acb4 dimers in complex with 3′3′-cGAMP and cartoon schematic illustrating mechanism of Acb4 tetramer assembly. (C) Surface electrostatic view of Acb4 binding pocket and polder omit map of 3′3′-cGAMP contoured at 2.9 σ. (D) EMSA analysis of Acb4 3′3′-cGAMP complex formation. A titration of recombinant Acb4 was incubated with 20 nM α^32^P-radiolabeled 3′3′-cGAMP, and reactions were analyzed by nondenaturing polyacrylamide gel electrophoresis. Data are representative of at least *n* = 3 independent experiments. (E) Quantification of EMSA analysis to determine Acb4 3′3′-cGAMP affinity. The fraction bound (bound intensity / total intensity) was calculated for each protein concentration and fit to a single binding model. Data represent the mean ± SD from *n =* 2 independent experiments.

The Acb4 3′3′-cGAMP-binding site is situated in a pocket formed between an extended β_3_–α_4_ loop region of one protomer and helices α_4_ and α_5_ from the opposite dimeric unit. The binding pocket is recessed deep in the Acb4 protomer interface and exhibits an overall strongly positive charge (Figure 3C). 3′3′-cGAMP ligands bind within each of the four equivalent interfaces creating a 4:4 Acb4–3′3′-cGAMP assembly (Figures 3A and 3C). Like other anti-CBASS and anti-Thoeris viral sponge proteins^21,27–30^, the Acb4 assembly maintains a 1:1 or greater protein to ligand binding site ratio suggesting that maximizing this ratio may be a key shared feature of viral sponge protein evolution. Acb4 binds 3′3′-cGAMP with high affinity (∼157 nM *K*_D_), consistent with the strong interactions required to sequester nucleotide immune signals and prevent downstream antiviral responses (Figures 3D and 3E).

Key structural features in Acb4 enable the selective recognition of nucleotide immune signals in anti-phage defense. The 3′3′-cGAMP ligand binding site is formed at the interface between two partnering protomers (denoted Acb4_A_ and Acb4_B_) and a third protomer from the opposite dimeric unit (Acb4_C_) (Figure 4A). Cooperation between the three Acb4 protomers creates a recessed binding site that envelops the host immune signal and coordinates 3′3′-cGAMP specificity through a combination of nucleobase- and phosphodiester linkage-specific contacts (Figures 4B and 4C). Notably, the phage SPO1 Acb4_A_–Acb4_B_ interface creates distinct adenine and guanine nucleobase pockets that allow sequence-specific recognition of 3′3′-cGAMP (Figures 4A and 4B). Recognition of guanine occurs through Acb4 residues K36_A_ and E49_B_ that coordinate sequence-specific contacts with the nucleobase Watson-Crick edge. Guanine base recognition is further stabilized by a π–π stacking interaction with Acb4 F48_B_ (Figures 4B and 4C). In the opposite pocket, adenine recognition occurs through peptide backbone interactions between Acb4 residue N63_A_ and the adenine Watson-Crick edge, and an additional network of van der Waals contacts formed by residues V56_A_, F62_A_, and I66_A_ that sterically restrict access of guanine nucleobases (Figures 4B and 4C). Contacts between Acb4 residues Y57_A_, T70_A_, and K81_C_ and the diphosphate backbone of 3′3′-cGAMP further stabilize ligand recognition (Figures 4B, 4C, and S6B). Mutagenesis of Acb4 binding pocket residues resulted in decreased 3′3′-cGAMP binding, demonstrating the importance of both nucleobase- and phosphodiester linkage-specific contacts in coordinating effective ligand sequestration (Figures 4D and S6A).

**Figure 4.**
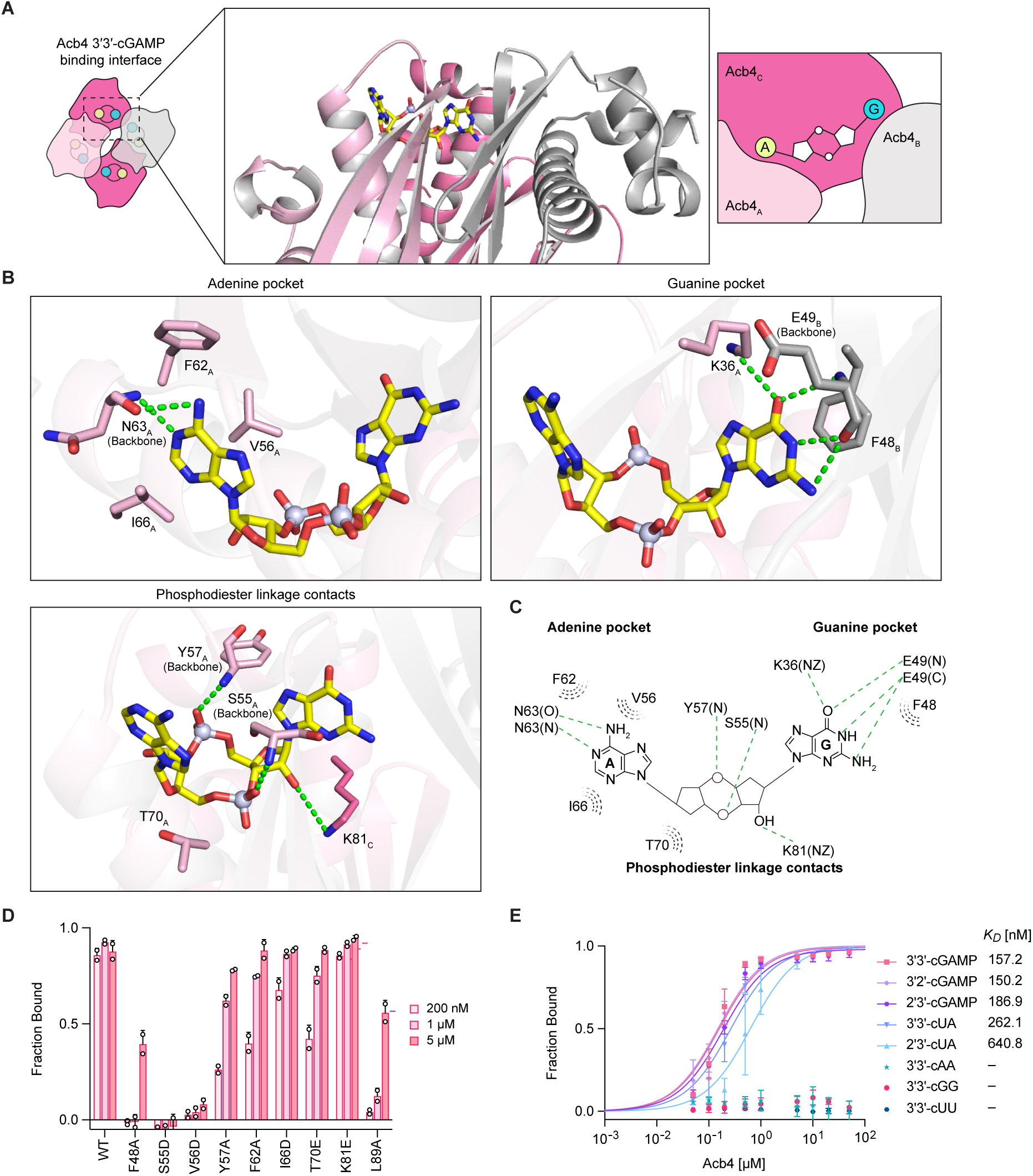
Structural basis of Acb4 nucleotide immune signal recognition. (A) Overview of the Acb4 ligand binding site with individual Acb4 protomers contacting 3′3′-cGAMP colored in light pink, magenta, and gray, respectively. Subscript denotes protomer chain of Acb4 tetramer. (B) Detailed view of Acb4 residues that interact the adenine base (top left), guanine base (top right), and phosphodiester linkages (bottom left) of 3′3′-cGAMP. Green dashed lines represent hydrogen bonding interactions and subscript denotes protomer chain of Acb4 tetramer. (C) Schematic highlighting key Acb4 residues that form contacts with 3′3′-cGAMP. Green dashed lines represent hydrogen bonds and grey dashed lines represent hydrophobic interactions. (D) Quantification of 3′3′-cGAMP binding with recombinant Acb4 proteins harboring point mutations in binding pocket residues. Data represent the mean ± SD from *n* = 2 independent experiments. For primary EMSA data, see Figure S6A. (E) Quantification of binding affinity of Acb4 with the cyclic dinucleotides 3′3′-cGAMP, 3′2′-cGAMP, 2′3′-cGAMP, 3′3′-cUA, 2′3′-cUA, 3′3′-cAA, 3′3′-cGG, and 3′3′-cUU calculated from EMSA analysis. Data represent the mean ± SD from *n* = 2 independent experiments. For primary EMSA data, refer to Figure S6C.

Nucleobase-specificity allows Acb4 to discriminate between CBASS signals from cyclic dinucleotides that regulate normal bacterial growth and intracellular signaling.^13,42,43^ No interaction was detected between Acb4 and either of the chemically related bacterial nucleotide second messengers 3′3′-c-di-AMP (3′3′-cAA) or 3′3′-c-di-GMP (3′3′-cGG), demonstrating that Acb4 does not bind cyclic dinucleotides with two guanine or two adenine bases (Figures 4E and S6C). We tested cGAMP isomers with alternative phosphodiester linkage backbones and observed that Acb4 also binds other CBASS signals including 2′3′-cGAMP and 3′2′-cGAMP with high affinity (*K*_D_ of ∼150 and ∼187 nM, respectively) (Figures 4E and S6C). Additionally, Acb4 recognizes the CBASS signals 3′3′-cUA and 2′3′-cUA, supporting that the guanine pocket can accommodate ligands with a smaller uridine nucleobase (Figures 4E and S6C). Together, these results define Acb4 nucleotide ligand specificity and explain the structural basis of inhibition of host CBASS anti-phage defense.

### Acb4 is the founding member of a widespread family of viral sponges

CBASS immunity spans all major bacterial phyla and is encoded in >16% of sequenced bacterial genomes.^10–13^ Given the prevalence of CBASS anti-phage defense, we hypothesized that other phages may also encode Acb4 homologs to inhibit bacterial immunity. We searched for SPO1 Acb4 homologs in the Pfam / InterPro database^44,45^ and identified 1,331 homologs that comprise a large family of uncharacterized proteins designated PF13876 (Table S3). Phylogenetic analyses of Acb4 revealed the presence of homologs in diverse phages that infect hosts spread across the major bacterial phyla Proteobacteria, Firmicutes, Bacteroidetes, and 38 unique genera (Figures 5A, S4B, and Table S3). Acb4 homologs are predominantly encoded in phage genomes or bacterial sequences with hallmark features of integrated prophage elements, suggesting the protein is predominantly encoded by viruses (Figure 5A).

**Figure 5.**
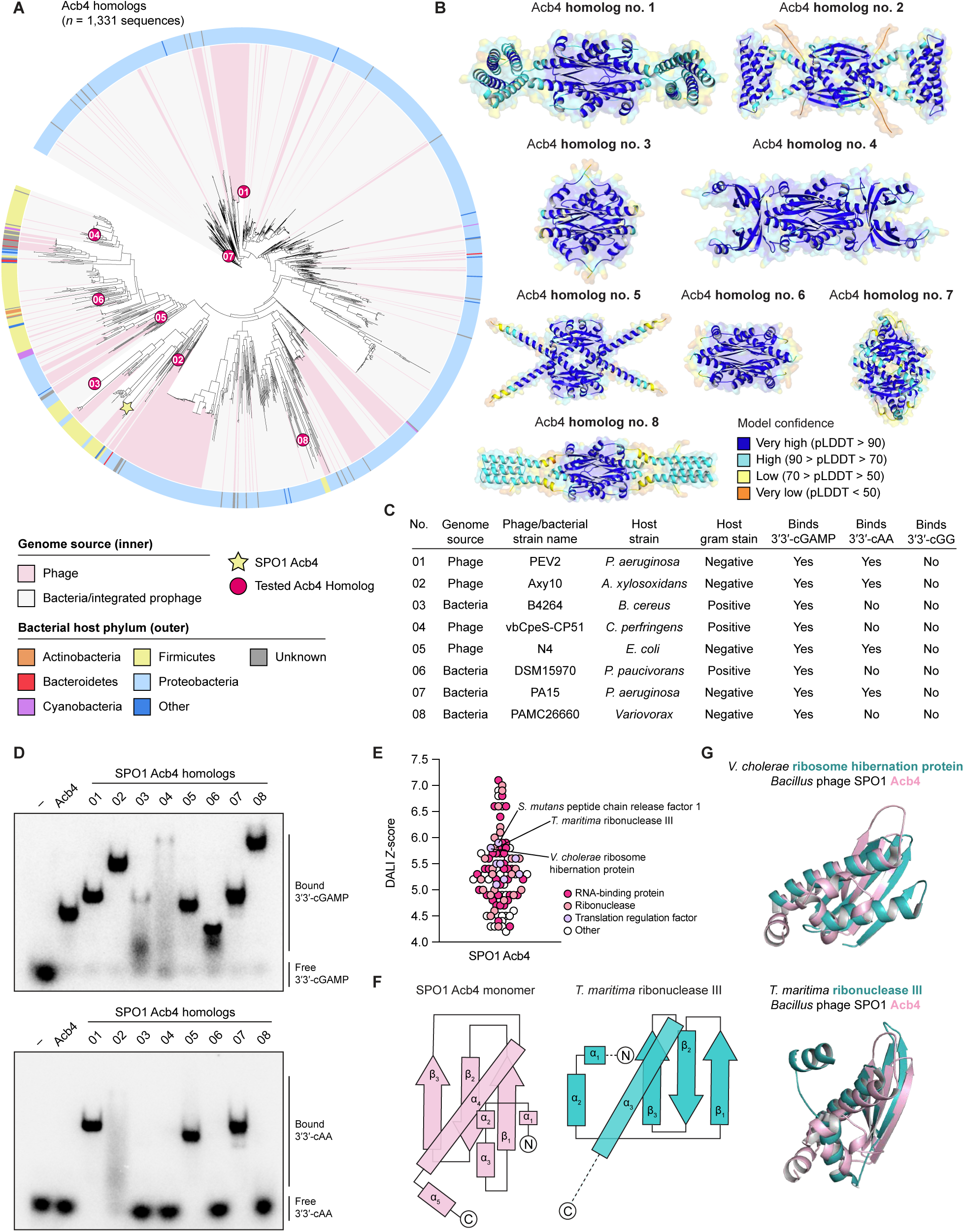
SPO1 Acb4 is the founding member of a widespread family of viral sponges. (A) Phylogenetic analysis of 1,331 Acb4 homologs. SPO1 Acb4 is denoted with a yellow star. Red circles with interior numbering represent Acb4 family proteins selected for bioinformatic and biochemical analysis. For additional information, refer to Figure S4 and Table S3. (B) Predicted structural models of Acb4 homologs generated using AlphaFold3 and colored by predicted local distance difference test (pLDDT). (C) Summary of purified Acb4 homologs and their binding properties. (D) Electrophoretic mobility shift assay to determine binding of SPO1 Acb4 and purified homologs against 3′3′-cGAMP (top gel) and 3′3′-cAA (bottom gel). Proteins were present at 5 µM and incubated with 20 nM ^32^P-labeled 3′3′-cGAMP or 3′3′-cAA, and bound complexes were visualized by nondenaturing polyacrylamide gel electrophoresis. Data are representative of at least *n* = 3 independent experiments. (E) DALI Z-scores for the top 100 hits for SPO1 Acb4 monomer when searched against all entries in the Protein Data Bank (PDB). (F) Topology map diagrams comparing *Bacillus* phage SPO1 Acb4 monomer with dsRNA-binding domain containing region of *Thermotoga maritima* ribonuclease III (PDB: 1O0W). Rectangles denote α-helices and arrows denote β-strands. (G) Structural comparison of SPO1 Acb4 with *Vibrio cholerae* ribosome hibernation factor (PDB: 4HEI) and *Thermotoga maritima* ribonuclease III (PDB: 1O0W). SPO1 Acb4 is colored in light pink and bacterial proteins are colored in teal.

To understand the diversity of Acb4 proteins, we selected eight Acb4 homologs for bioinformatic and functional characterization. We began by comparing the crystal structure of the *Bacillus* phage SPO1 Acb4 complex with predicted structural models of each Acb4 homolog generated using AlphaFold3 (Figures 5B and 5C).^46^ Consistent with the oligomeric assembly of SPO1 Acb4, all eight homologs folded with high confidence as tetramers, supporting that the core architecture of Acb4 is conserved throughout this family of viral sponge proteins (Figure 5B). Interestingly, although the core SPO1 Acb4 protein architecture is conserved across all eight homologs, we observed that several Acb4 family proteins harbored additional extensions, suggesting that select phage Acb4 homologs may be adapted for additional functions (Figure 5B).

We next cloned and recombinantly expressed all eight Acb4 homologs as purified proteins (Figures 5A–C and S4A). All eight Acb4 homologs bound 3′3′-cGAMP, supporting a broad role for Acb4 in inhibiting CBASS defense (Figures 5C and 5D). We measured the ability of Acb4 homologs to selectively bind nucleotide immune signals and avoid recognition of the cellular signals 3′3′-cGG and 3′3′-cAA. Most bacteria synthesize the cyclic dinucleotide 3′3′-cGG as a second messenger that regulates diverse processes including biofilm formation, cell cycle progression, and virulence.^42,47,48^ Gram-positive bacteria additionally synthesize 3′3′-cAA as a nucleotide signal that regulates cell wall homeostasis, host transcription, and ion transport.^43,49,50^ Consistent with this functional distribution of signaling roles in bacteria, no Acb4 homologs in our panel recognized the cyclic dinucleotide 3′3′-cGG (Figures 5C and S4C). Strikingly, we observed a clear pattern where only Acb4 homologs from phages that infect Gram-negative bacteria hosts are capable of binding 3′3′-cAA (Figures 5C and 5D). These results suggest that viral sponges likely evolve to adapt to the local host signaling environment to exclusively target nucleotide signals in anti-phage defense.

To explore the potential evolutionary origin of immune evasion proteins, we used Dali^51^ to identify proteins with predicted structural homology to the SPO1 Acb4 monomer (Figure 5E). Surprisingly, our analyses revealed strong structural homology between Acb4 and bacterial proteins with a double-stranded RNA-binding domain (dsRBD) (Figures 5E and 5F). dsRBD-containing proteins are highly conserved RNA-binding proteins across all kingdoms of life that are defined by an α–β–β–β–α domain topology that spans approximately 70 amino acids.^52–54^ Top bacterial hits with structural homology to Acb4 included diverse ribonucleases and proteins that regulate host translation, including peptide chain release factors and ribosome hibernation proteins (Figures 5E–5G). These results support a potential model of Acb4 evolution where a common RNA-binding domain was adapted into a viral sponge by oligomerization and creation of ligand-binding pockets for nucleotide immune signal recognition. Together, our findings reveal Acb4 as a widespread family of viral sponge proteins that have been adapted to selectively inhibit nucleotide immune signaling and enable phage immune evasion (Figure 6A).

**Figure 6.**
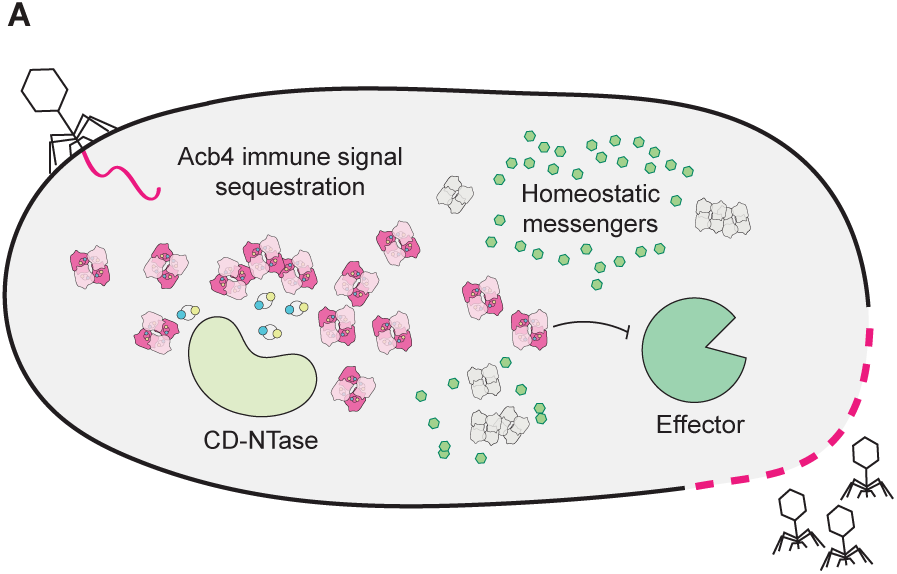
Summary model of CBASS immune evasion by Acb4. (A) Conceptual model of Acb4-mediated inhibition of CBASS anti-phage defense. During phage infection, Acb4 inhibits CBASS signaling by sequestering nucleotide immune signals while avoiding disruption of signals required for normal cellular homeostasis.

## Discussion

In this study, we discover Acb4 from the *Bacillus* phage SPO1 as the founding member of a widespread family of >1,300 anti-CBASS immune evasion proteins. We demonstrate that Acb4 is a phage-encoded sponge that sequesters CBASS signals to subvert host anti-phage defense, highlighting viral inhibition of host nucleotide immune signals as a conserved mechanism of immune evasion.^1,2^ Our results define the mechanism and structural basis for how Acb4 discriminates between immune signals and other cellular signals, explaining how viral sponge proteins evolve to selectively antagonize host immunity.

CBASS immunity is one of the most common forms of bacterial anti-phage defense^10–13^, creating strong evolutionary pressure for phages to evolve mechanisms of immune evasion. Our forward biochemical screen of phage-infected lysates identifies Acb4 as a viral sponge protein that binds CBASS nucleotide immune signals and prevents downstream Cap effector protein activation (Figures 1 and 2). CBASS anti-phage defense operons are exceptionally diverse, with >5,000 known CD-NTase enzymes that synthesize distinct classes of nucleotide immune signals and dozens of families of Cap effector proteins.^4,13,14^ To overcome this diversity, we show that phage Acb4 effectively targets a range of CBASS nucleotide immune signals including 3′3′-cGAMP, 2′3′-cGAMP, 3′2′-cGAMP, 3′3′-cUA, and 2′3′-cUA (Figures 1 and 4). *In vivo*, we use chimeric phage T4 variants to demonstrate that Acb4 expression is sufficient to inhibit *Yersinia aleksiciae* CBASS 3′3′-cUA immune signaling and restore phage replication (Figure 2). Interestingly, although SPO1 Acb4 is capable of binding 3′2′-cGAMP *in vitro*, we observed that a T4 phage expressing SPO1 Acb4 is unable to overcome 3′2′-cGAMP signaling *in vivo* during *Citrobacter portucalensis / Escherichia coli* CBASS immunity (Figure 2). In contrast, T4 phages expressing the nuclease Acb1 that only weakly cleaves 3′2′-cGAMP could overcome *Citrobacter portucalensis / Escherichia coli* CBASS immunity, suggesting that phages may evolve to encode multiple Acb proteins due to distinct advantages of sponge and nuclease immune evasion proteins against specific CBASS operons.

Effective CBASS evasion requires viral sponge proteins that precisely recognize nucleotide immune signals while avoiding non-specific interactions with closely related signals that control bacterial homeostasis. Our structural and biochemical analyses of *Bacillus* phage SPO1 Acb4 and additional Acb4 homologs define key molecular features that enable ligand specificity. To sequester nucleotide immune signals, SPO1 Acb4 and all known viral sponge proteins assemble into oligomeric complexes with ligand binding pockets formed at each protomer-protomer interface (Figure 3). In Acb4, the ligand binding site is formed through contacts donated by three separate protomers, creating discrete nucleotide pockets that allow specific recognition of the guanine and adenine bases in common CBASS signals like 3′3′-cGAMP (Figure 4). In addition to immune signaling, bacteria use 3′3′-cGG and 3′3′-cAA cyclic dinucleotides to regulate cellular stress, biofilm formation, cell motility, and virulence.^43,55,56^ Gram-positive bacteria like *Bacillus subtilis* synthesize both 3′3′-cGG and 3′3′-cAA signals, while Gram-negative bacteria like *E. coli* synthesize only 3′3′-cGG.^43,55^ Strikingly, our biochemical studies demonstrate that Acb4 ligand-specificity closely follows this phylogenetic pattern to avoid targeting host bacteria homeostatic signals. Acb4 homologs from phages infecting Gram-positive hosts avoid both 3′3′-cGG and 3′3′-cAA recognition, while in contrast, Acb4 homologs from phages infecting Gram-negative hosts avoid only 3′3′-cGG (Figures 4 and 5). We hypothesize that advantages of viral sponge proteins adapting to specific host cellular signaling environments include limiting competition with abundant homeostatic signals that may saturate ligand-binding sites and avoiding disruption of host signaling pathways that may reduce host cell fitness.

Finally, our structural and phylogenetic analyses of the Acb4 immune evasion protein family reveal insight into the potential evolutionary origin of anti-CBASS proteins. Structure-guided homology searches show that Acb4 monomers share architectural homology with the dsRBM domain shared in bacterial RNA-binding proteins, including ribonucleases, peptide chain release factors, and small RNA interacting proteins, suggesting that Acb4 may have originated from a common bacterial RNA-binding fold (Figure 5). The anti-CBASS nuclease protein Acb1 also shares structural homology with bacterial host proteins belonging to the 2H phosphoesterase superfamily^21,25,26^, supporting that a common route of Acb protein evolution may be acquisition and co-option of host proteins for subversion of immune defense. Acb4 joins a growing list of recently discovered viral sponge proteins including Tad1, Tad2, and Acb2 that sequester nucleotide immune signals to inactivate host anti-phage defense.^21,27–30^ Our results expand viral sponges as a major class of immune evasion proteins and explain the molecular principles by which phages adapt to specifically inhibit host antiviral immunity.

## Supporting information

Chang et al Table S1

Chang et al Table S2

Chang et al Table S3

## Acknowledgements

We are grateful to Y. Li and members of the Kranzusch lab for helpful comments and discussion. Mass spectrometry was performed at the Taplin Mass Spectrometry Facility at Harvard Medical School with assistance from R. Tomaino. The work was funded by grants to P.J.K. from the Pew Biomedical Scholars program, the Burroughs Wellcome Fund PATH program, The Mathers Foundation, The Mark Foundation for Cancer Research, the Cancer Research Institute, and the National Institutes of Health (1DP2GM146250-01), grants to E.T.C from the Claudia Adams Barr Program, the Lavine Family Fund, the Pew Charitable Trust, NIH DK123095, NIH AG071966, NIH AI175317, The Smith Family Foundation, and the American Federation for Aging Research, and grants to R.S. from the European Research Council (ERC-AdG GA 101018520), the Israel Science Foundation (MAPATS Grant 2720/22), the Ernest and Bonnie Beutler Research Program of Excellence in Genomic Medicine, the Deutsche Forschungsgemeinschaft (SPP 2330, grant 464312965), the Minerva Foundation with funding from the Federal German Ministry for Education and Research, the Institute for Environmental Sustainability (IES) and the Center for Immunotherapy at the Weizmann Institute of Science, a research grant from the Estate of Marjorie Plesset, and the Knell Family Center for Microbiology. R.B.C. is supported through a Landry Cancer Biology Research Fellowship (Harvard Faculty of Arts and Sciences), S.J.H. is supported through a Cancer Research Institute Irvington Postdoctoral Fellowship (CRI3996), D.R.-B. is supported through the National Science Foundation Graduate Research Fellowship Program, and N.B. is supported by the Deutsche Forschungsgemeinschaft (DFG: Projektnummer 501493132). X-ray data were collected at The Center for Bio-Molecular Structure (CBMS) that is primarily supported by the NIH-NIGMS through a Center Core P30 Grant (P30GM133893), and by the DOE Office of Biological and Environmental Research (KP1607011). NSLS2 is a U.S.DOE Office of Science User Facility operated under Contract No. DE-SC0012704. This publication resulted from the data collected using the beamtime obtained through NECAT BAG proposal # 311950.

## Author Contributions

Experiments were designed and conceived by R.B.C. and P.J.K. Biochemical screen was performed by R.B.C. with assistance from H.C.T. using phage-infected lysates prepared by T.W. and R.S. Thermal proteome profiling experiments and analyses were performed by N.B. and E.T.C. Phage Acb4 identification and biochemical experiments were performed by R.B.C. with assistance from H.C.T. Engineering of recombinant phages was performed by S.J.H. and D.R.*-*B. Phage challenge experiments were performed by R.B.C. Crystallography experiments, structural analyses, and bioinformatic analyses were performed by R.B.C. The manuscript was written by R.B.C. and P.J.K. All authors contributed to editing the manuscript and support the conclusions.

## SI Figure Legends

**Figure S1.**
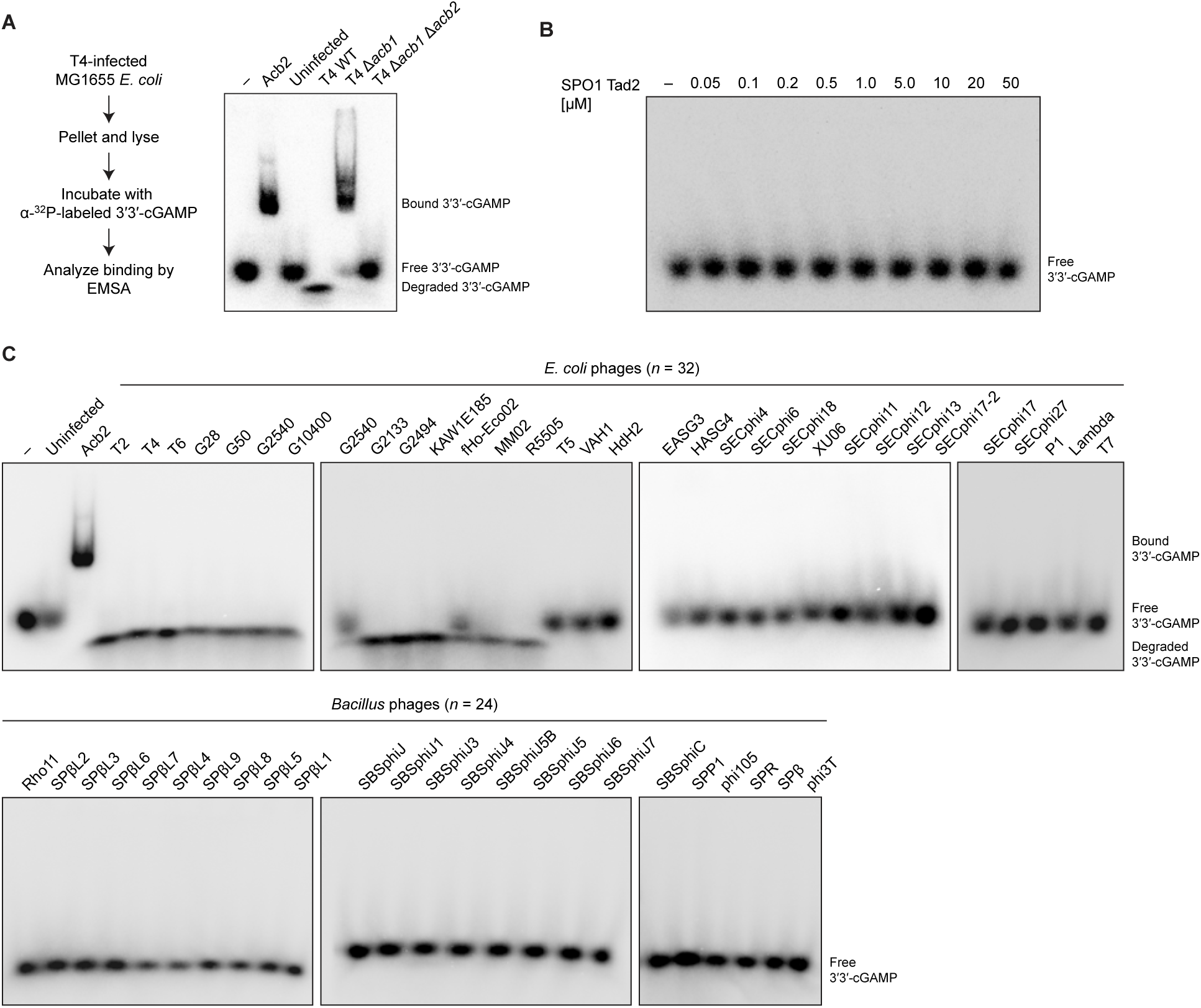
Biochemical screen of 3′3′-cGAMP binding activity in *Bacillus subtilis* and *Escherichia coli* phage-infected lysates, related to Figure 1. (A) Schematic of screening approach and EMSA analysis of 3′3′-cGAMP binding and degradation activity detected in wildtype and recombinant T4 phage-infected lysates. Data are representative of at least *n* = 3 independent experiments. (B) EMSA analysis of the binding of recombinant SPO1 Tad2 and 3′3′-cGAMP. Data are representative of *n* = 2 independent experiments. (C) Primary data from a biochemical screen for 3′3′-cGAMP binding activity in 32 *Escherichia coli* phages and 24 *Bacillus subtilis* phages. Phage-infected lysates were incubated with α^32^P-radiolabeled 3′3′-cGAMP, and binding or ligand degradation activity was visualized by EMSA. Data are representative of *n* = 2 independent experiments. For list of phages used in biochemical screen, refer to Table S1.

**Figure S2.**
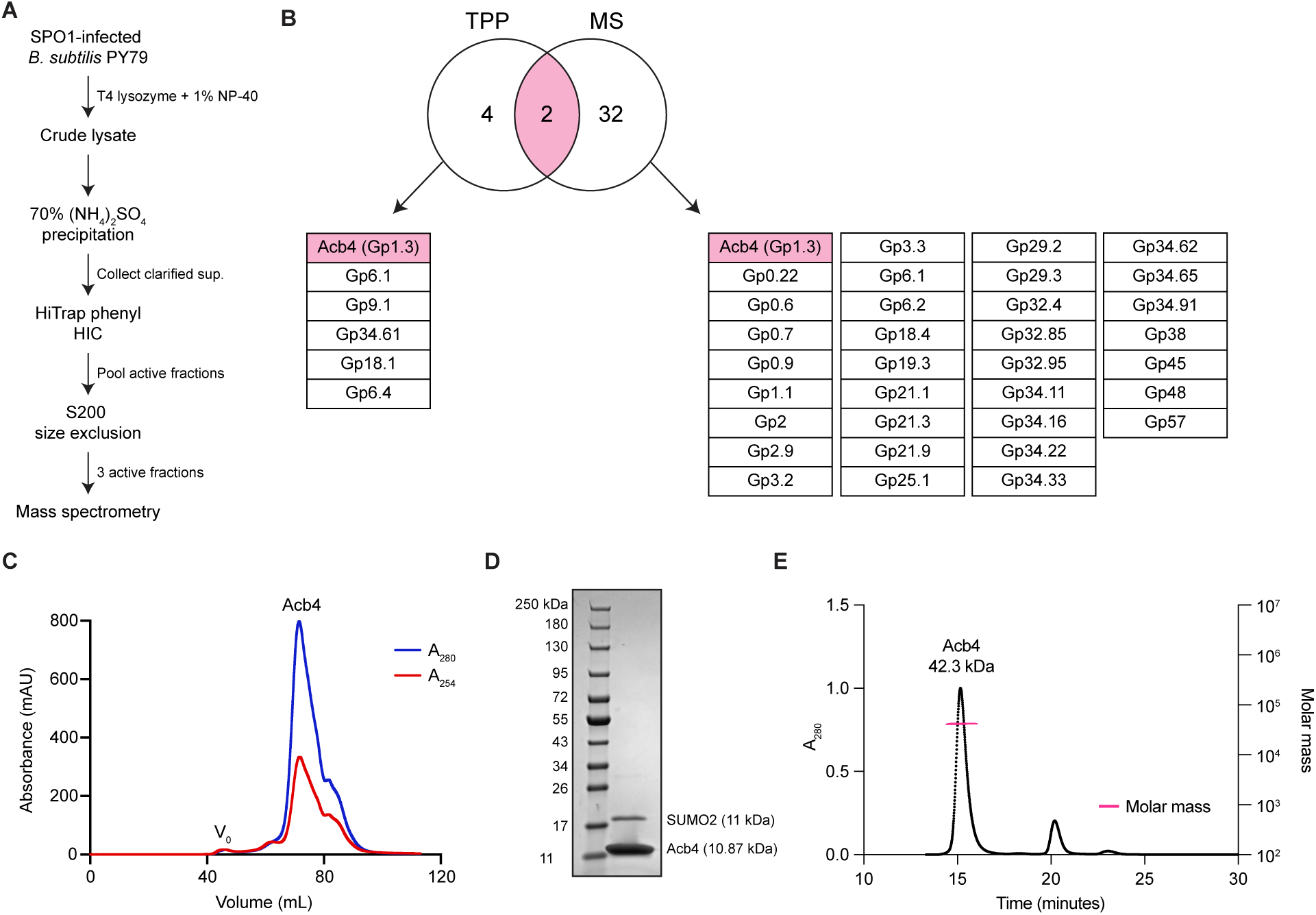
Identification and purification of *Bacillus* phage SPO1 Acb4, related to Figure 1. (A) Schematic of biochemical fractionation strategy used to enrich 3′3′-cGAMP binding activity from crude *Bacillus* phage SPO1-infected lysate. (B) Venn diagram comparing candidate Acb4 proteins identified independently through thermal proteome profiling (TPP) and biochemical fractionation coupled with mass spectrometry (MS). (C) Purification of phage SPO1 Acb4 from *Escherichia coli*. SPO1 Acb4 was expressed as an N-terminal 6×His-SUMO fusion protein and purified by Ni-NTA affinity chromatography followed by S75 size-exclusion chromatography. (D) SDS-PAGE analysis of recombinant *Bacillus* phage SPO1 Acb4 visualized by Coomassie blue staining. (E) Size-exclusion chromatography with multi-angled light scattering analysis of recombinant *Bacillus* phage SPO1 Acb4. The Acb4 complex migrates at ∼42.3 kDa, consistent with a tetrameric assembly.

**Figure S3.**
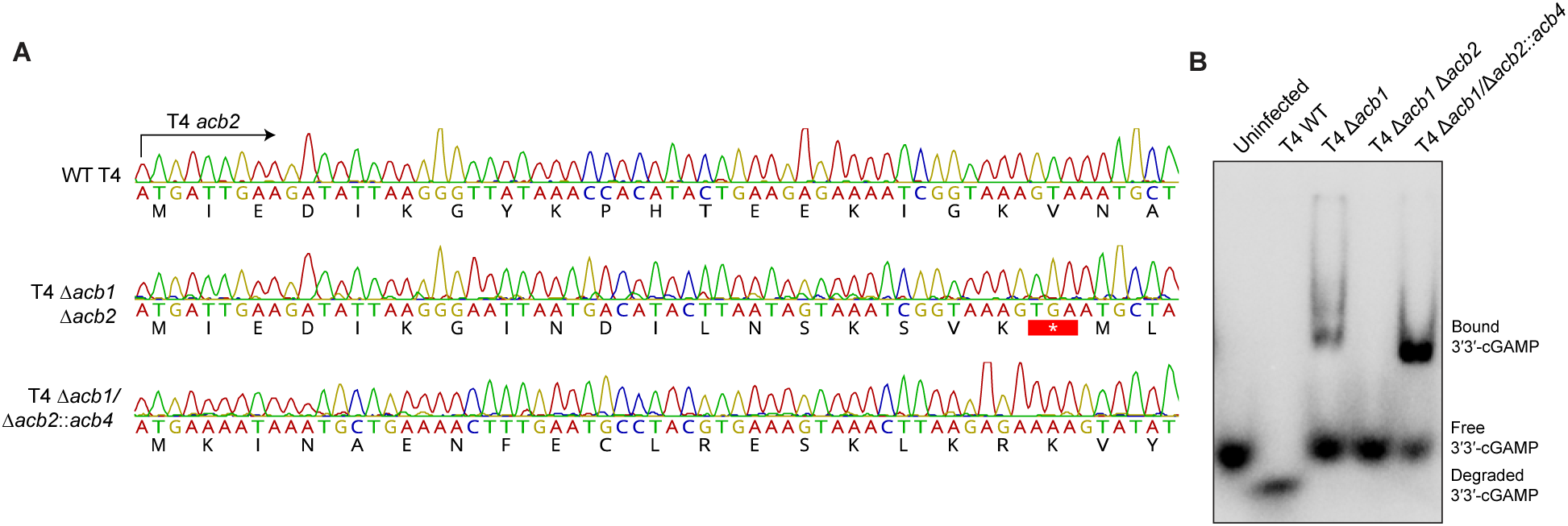
Generation and validation of mutant phage T4 viruses, related to Figure 2. (A) Sanger sequencing reads confirming successful replacement of endogenous phage T4 *acb2* with *Bacillus* phage SPO1 *acb4*. (B) EMSA analysis demonstrating 3′3′-cGAMP cleavage and binding activity in wildtype and recombinant phage T4 viruses.

**Figure S4.**
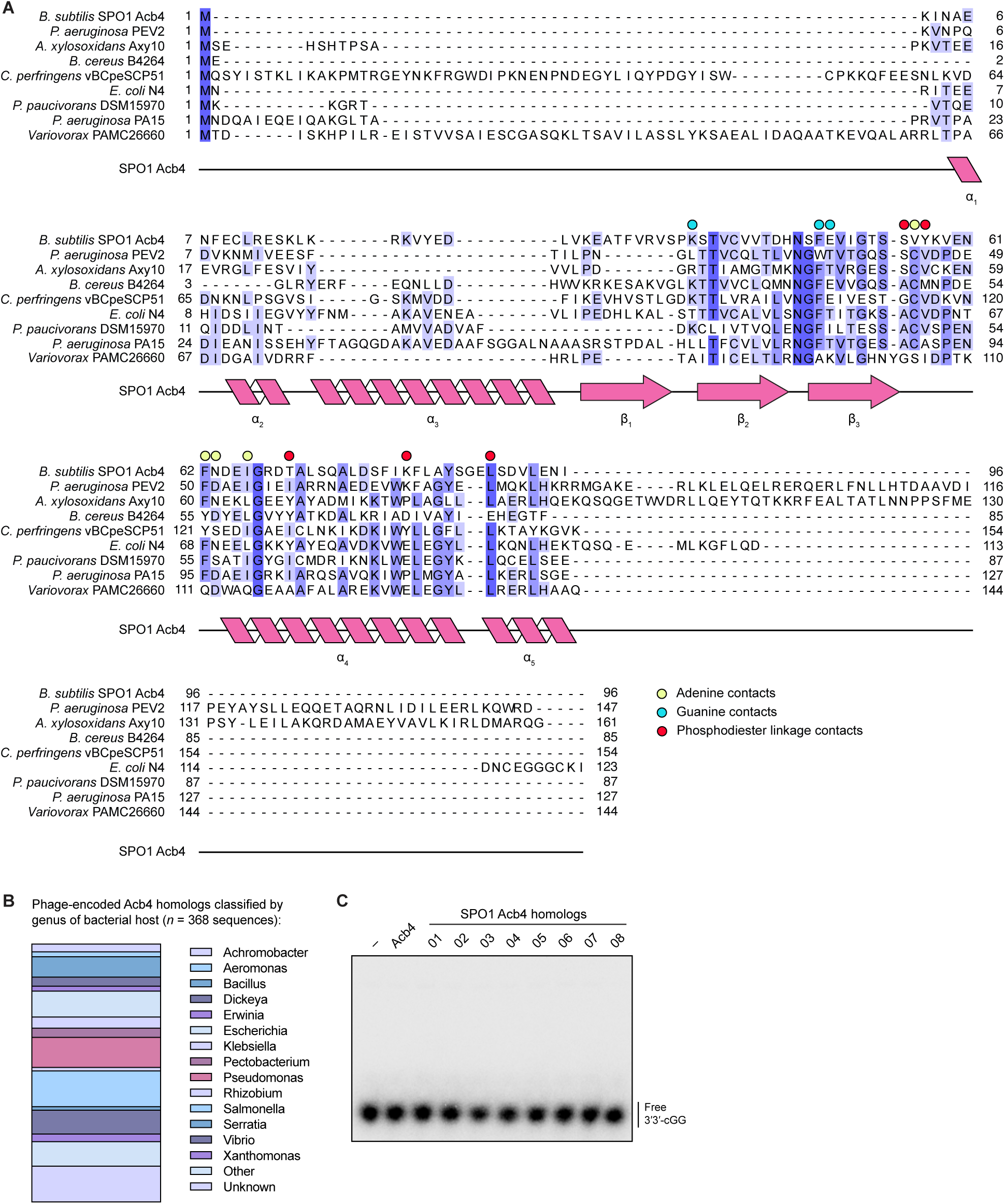
Acb4 structural characterization and diversity of Acb4 homologs, related to Figures 3–5. (A) Sequence alignment of *Bacillus* phage SPO1 Acb4 and Acb4 homologs. (B) Bar graph illustrating distribution of phage-encoded Acb4 homologs classified by genus of bacterial host. (C) EMSA analysis to determine binding of SPO1 Acb4 and purified homologs against the bacterial second messenger 3′3′-cGG. Proteins were present at 5 µM and incubated with 20 nM ^32^P-labeled 3′3′-cGG, and reactions were visualized by nondenaturing polyacrylamide gel electrophoresis. Data are representative of at least *n* = 3 independent experiments.

**Figure S5.**
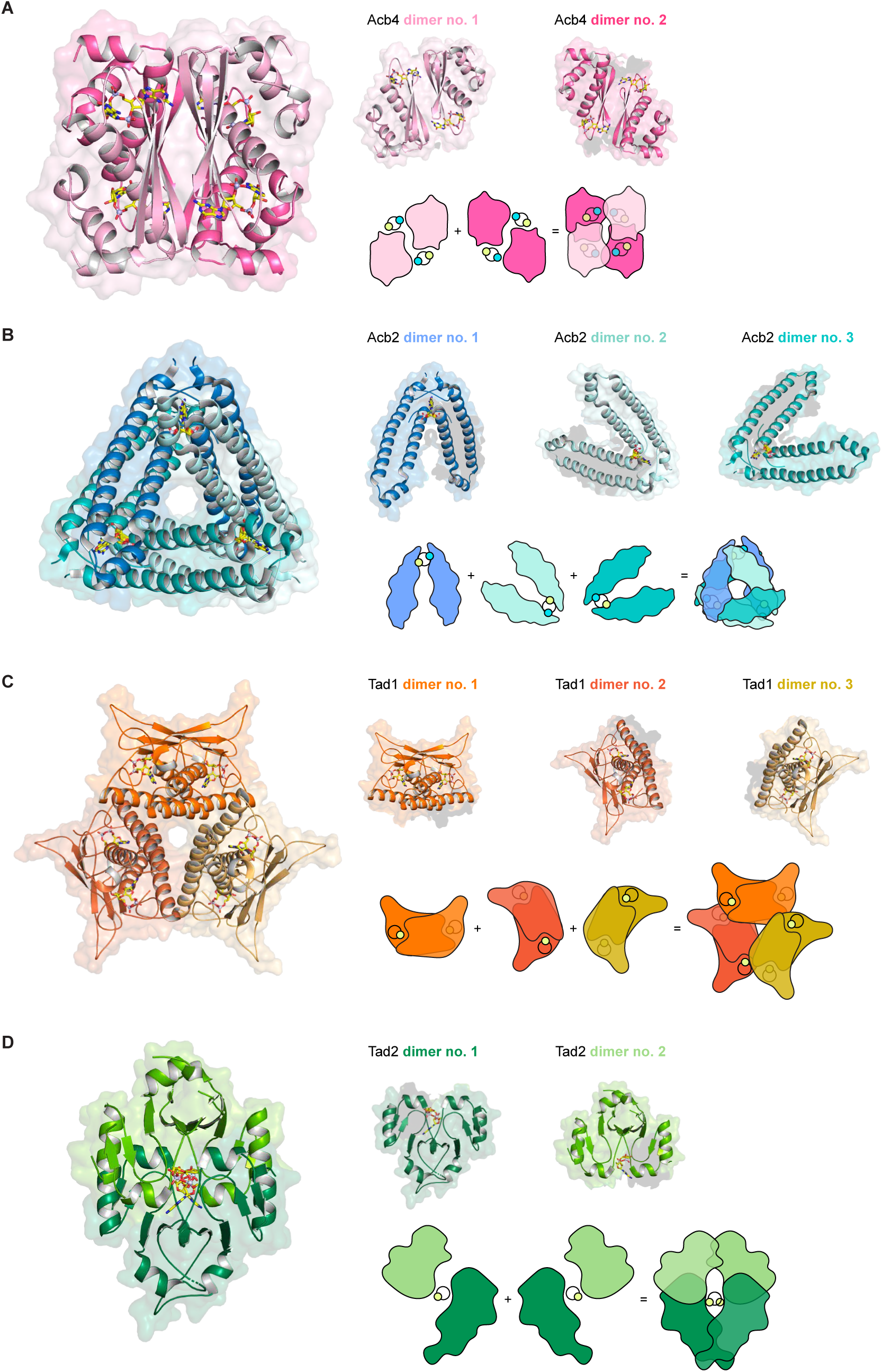
Structural comparison of *Bacillus* phage SPO1 Acb4 with viral sponge families, related to Figures 3 and 4. (A) Overview of *Bacillus* phage SPO1 Acb4 tetramer in complex with 3′3′-cGAMP (PDB: 9E4W) and cartoon illustrating Acb4 assembly. (B) Overview of *Pseudomonas* phage PaP2 Acb2 hexamer in complex with 3′3′-cGAMP (PDB: 8H2J) and cartoon illustrating Acb2 assembly. (C) Overview of *Clostridium botulinum* prophage Tad1 hexamer in complex with 1′′–3′ gcADPR (PDB: 8SMD) and cartoon illustrating Tad1 assembly. (D) Overview of *Bacillus* phage SPO1 Tad 2 tetramer in complex with 1′′–3′ gcADPR (PDB: 8SMF) and cartoon illustrating Tad2 assembly.

**Figure S6.**
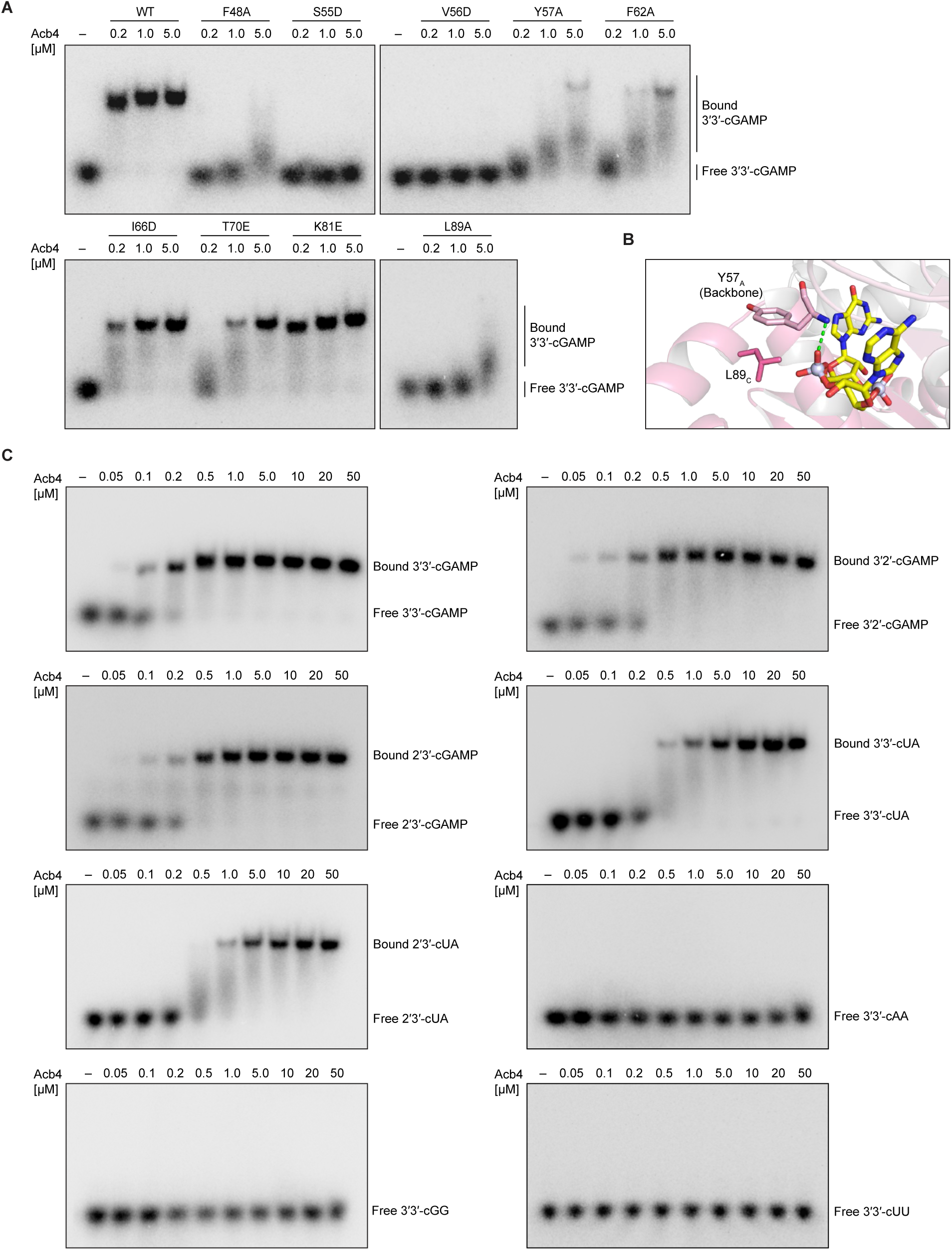
Biochemical and mutagenesis analysis of Acb4 ligand interaction, related to Figure 4. (A) Primary EMSA analysis data measuring 3′3′-cGAMP complex formation with Acb4 nucleotide binding pocket point mutants in Figure 4D. Data are representative of *n* = 2 independent experiments. (B) Detailed view of 3′3′-cGAMP phosphodiester backbone and stabilization by Acb4 residues Y57_A_ and L89_C_. Green dashed lines represent hydrogen bonding interactions and subscript denotes protomer chain of Acb4 tetramer. (C) Primary EMSA analysis data for quantification of binding affinity in Figure 4E of Acb4 with 3′3′-cGAMP, 3′2′-cGAMP, 2′3′-cGAMP, 3′3′-cUA, 2′3′-cUA, 3′3′-cAA, 3′3′-cGG, and 3′3′-cUU. Data are representative of *n* = 2 independent experiments.

**Table S1. Summary of phages and lysate sampling times used in biochemical screen, related to Figure 1**

**Table S2. Crystallographic Statistics, related to Figures 3 and 4**

**Table S3. Summary of Acb4 homolog sequences and corresponding taxonomic information, related to Figure 4**

## STAR METHODS

### RESOURCE AVAILABILITY

#### Lead Contact

Further information and requests for resources and reagents should be directed to and will be fulfilled by the Lead Contact, Philip Kranzusch.

#### Materials Availability

This study did not generate new unique reagents.

#### Data and Code Availability

- Coordinates of *Bacillus* phage SPO1 Acb4 in complex with 3′3′-cGAMP are publicly available in the Protein Data Bank under the accession number 9E4W.
- This paper does not report original code.
- Any additional information required to reanalyze the data reported in this paper is available from the lead contact upon request.

### METHOD DETAILS

#### Preparation of phage-infected lysates in biochemical screen

Overnight cultures of *E. coli* MG1655 (ATCC 47076) and *B. subtilis* BEST7003^57^ were diluted 1:100 in 250 mL MMB media and grown at 37°C and 30°C, respectively. Back-diluted cultures were infected with phages at an OD_600_ of ∼0.30 (early log phase) at a final multiplicity of infection of 2. Following phage infection, a 5 mL aliquot of the infected culture was harvested prior to phage-induced collapse by centrifugation at 3,200 × g for 5 minutes at 4°C. The infected pellets were flash frozen in dry ice and ethanol. Phage-infected *E. coli* pellets were resuspended in 250 μL of bacterial lysis buffer (20 mM HEPES-KOH pH 7.5, 150 mM NaCl, 5 mM MgCl_2_, 1 mM MnCl_2_, 10% glycerol, 1% NP-40, and 1 mM DTT) and incubated for 30 minutes at room temperature. Phage-infected *Bacillus* pellets were resuspended in PBS and pre-treated with 1 mg mL^−1^ of T4 lysozyme (ThermoFisher) at 37°C for 10 minutes, followed by treatment with 400 μL of bacterial lysis buffer and incubation at room temperature for 30 minutes. Lysates were clarified by centrifugation at 20,000 × g for 5 minutes and the resultant supernatant was flash frozen in liquid nitrogen and stored at −80°C. A summary of the *E. coli* and *Bacillus* phages included in this study and the respective sampling times is provided in Table S1.

#### Synthesis of radiolabeled cyclic nucleotide immune signals

Cyclic nucleotides used for binding analyses were synthesized using the following purified recombinant enzymes: *Vibrio cholerae* DncV^4^: 3′3′-cAA, 3′3′-cGAMP, and 3′3′-cGG; human cGAS^58^: 2′3′-cGAMP; *Drosophila eugracilis* cGLR1^59^: 3′2′-cGAMP; *Crassostrea virginica* cGLR1^60^: 2′3′-cUA; *Yersinia aleksiciae* CdnE^19^: 3′3′-cUA and 3′3′-cUU. Cyclic nucleotide synthesis was performed at 37°C for ∼20 hours in 10 μL reactions containing a final concentration of 5 µM recombinant enzyme, 20 µM appropriate nucleotide triphosphates (NTPs), trace amounts of appropriate α-^32^P-labeled NTP (Revvity), 25 mM KCl, 5 mM MgCl_2_, 1 mM MnCl_2_, 1 mM TCEP and 50 mM Tris-HCl pH 7.5 (DncV, human cGAS, *Drosophila eugracilis* cGLR1, *Crassostrea virginica* cGLR1) or pH 9.0 (*Ya*CdnE). Following overnight incubation, reactions were centrifuged at 17,000 × g for 1 minute to remove any precipitated protein. To digest unincorporated NTPs, clarified synthesis reactions were treated with 2 μL of Quick CIP (NEB) followed by incubation for ∼30 minutes at 37°C and heat inactivation for 2 minutes at 80°C. To confirm cyclic nucleotide synthesis, ∼1 μL of the final reaction product was spotted on a PEI cellulose thin-layer chromatography plate (Sigma Aldrich) and developed in a 1.5 M KH_2_PO_4_ (pH 3.8) buffer for 90 minutes. Plates were dried at room temperature for ∼20 minutes, exposed to a storage phosphor screen, and imaged with a Typhoon Trio Variable Mode Imager System (GE Healthcare).

#### Electrophoretic mobility shift assay

Electrophoretic mobility shift assays were used to assess nucleotide immune signal binding activity with either recombinant protein or phage-infected lysates. Recombinant proteins were diluted to concentrations ranging from 250 nM to 50 µM in a reaction buffer containing 50 mM Tris-HCl pH 7.5, 5 mM MgCl_2_, 10 mM KCl, and 1 mM TCEP. Diluted proteins or phage-infected lysates were incubated with 20 nM α^32^P-radiolabeled cyclic nucleotide for 30 minutes on ice. Reactions were resolved on a 7.2-cm nondenaturing 6% polyacrylamide gel run at 100V for 50 minutes in 0.5× TBE buffer. The gel was fixed in a solution containing 40% ethanol and 10% acetic acid and dried at 80°C for 30 minutes. The dried gel was exposed to a storage phosphor screen overnight and visualization was performed using a Typhoon Trio Variable Mode Imager System (GE Healthcare). The intensity of the radiolabeled signal was quantified using ImageQuant 5.2 software.

#### SPO1 Acb4 activity-guided fractionation and mass spectrometry analysis

An overnight culture of *B. subtilis* PY79^61^ was diluted 1:50 in a total volume of 1 L LB media and grown at 37°C with shaking (230 rpm). Cultures were grown to an OD_600_ of ∼0.30–0.40 (early log phase) and infected with phage SPO1 at a final multiplicity of infection of 2. Infected cultures were pelleted ∼70 minutes post-infection by centrifugation at 7,000 × g for 20 minutes. SPO1-infected pellets were resuspended in 10 mL of 50 mM Tris-HCl pH 7.5 and treated with 50 μL of T4 lysozyme (NEB) for 15 minutes at room temperature with gentle shaking. Lysates were further treated with 14 mL of bacterial lysis buffer (20 mM HEPES-KOH pH 7.5, 150 mM NaCl, 5 mM MgCl_2_, 1 mM MnCl_2_, 10% glycerol, 1% NP-40, and 1 mM DTT) for 30 minutes on ice with intermittent vortexing. Lysates were centrifuged at 74,000 × g for 30 minutes and the supernatant was clarified by 0.22 µm filtration. SPO1-infected lysates were subjected to a 70% (NH_4_)_2_SO_4_ cut for 1 hour at 4°C with continuous stirring, then centrifuged at 74,000 × g for 20 minutes to remove precipitated proteins. The clarified cut supernatant was passed through a 0.22 µm filter and fractionated by hydrophobic interaction chromatography using a 5 mL HiTrap Phenyl column (Cytiva) then eluted with a gradient of 2.00 M–0.00 M (NH_4_)_2_SO_4_. Active hydrophobic interaction chromatography fractions were pooled, concentrated, and further fractionated by size-exclusion chromatography with a Superdex 200 Increase 10/300 GL column (Cytiva). An active fraction containing peak 3′3′-cGAMP binding activity was selected for analysis by label-free mass spectrometry to identify candidate SPO1 genes. Acb4 was identified as the product of the SPO1 gene *1.3*.

#### Thermal proteome profiling

##### Sample preparation and protein precipitation

SPO1-infected phage lysates were prepared as described above in a lysis buffer containing 20 mM HEPES-KOH pH 7.5, 150 mM NaCl, 5 mM MgCl_2_, 1 mM MnCl_2_, 10% glycerol, and 1 mM DTT. Phage-infected lysates were quantified using the Quick Start Bradford Protein Assay (BioRad), diluted in lysis buffer to a final concentration of 2 mg mL^−1^, and flash frozen in liquid nitrogen. The SPO1-infected lysate (200 μL at 2 mg mL^−1^) was thawed on ice and combined with 200 μL of lysis buffer for control samples or buffer supplemented with 200 µM synthetic 3′3′-cGAMP or 3′3′-cGG (Biolog Life Science Institute). Lysates were incubated for 15 minutes at room temperature and heated for 3 minutes across a temperature gradient of 48–58°C in a PCR cycler. Samples were equilibrated to 21°C and centrifuged for 90 seconds at 21,000 × g. The supernatant was transferred to a fresh tube and proteins were denatured and reduced with 1% sodium dodecyl sulfate (SDS) and 5 mM tris(2-carboxyethyl)phosphine (TCEP) for 15 minutes at 37°C and subsequently labeled with 20 mM iodoacetamide. Labeled proteins were precipitated on ice using 3× volume methanol, 1× volume chloroform, and 2.5× volume water, then centrifuged at 15,000 × g for 15 minutes at 4°C. The resulting protein precipitate was washed twice with 1 mL methanol, followed by centrifugation at 15,000 × g for 5 minutes at 4°C.

##### Protein digestion and TMT labeling

Precipitated protein pellets (∼100 µg protein) were airdried to evaporate methanol and resuspended in 100 μL 200 mM N-(2-Hydroxyethyl)piperazine-N′-(3-propanesulfonic acid) (EPPS) pH 8.0 supplemented with trypsin (Promega) and LysC (Wako). Protein lysates were digested overnight with vigorous shaking at 37°C. Following overnight incubation, samples were centrifuged for 10 min at 21,000 × g, and peptide concentration was determined by microBCA (Thermo Scientific). Equal amounts of peptides for each sample (50 µg) were transferred into fresh tubes and samples were brought to a final volume of 100 μL in 200 mM EPPS pH 8.0. Peptides were labeled with 10 µl of TMTpro (16-plex) reagents in acetonitrile for 1 hour in the dark at RT and vortexed intermittently. Following TMT-labeling, 2 µl of each sample was combined into 140 μL 1% formic acid, desalted, and analyzed via LC-MS. Upon completion of the ratio check analysis, the labeling reaction was quenched by addition of 5 µl of 5% hydroxylamine and incubation for 15 min at RT. Samples were pooled at equal amounts according to the ratio check and diluted with 12 mL 1% FA in H_2_O and subjected to gravity flow-driven C18 solid-phase extraction (200 mg Sep-Pak, Waters) and vacuum dried. Samples were fractionated into 6 fractions using a Pierce^TM^ High pH Reversed-Phase Peptide Fractionation Kit (Thermo Scientific), followed by desalting via stage tip.

##### LC-MS/MS parameters

An Orbitrap Eclipse Tribrid Mass Spectrometer (Thermo) coupled with an Easy-nLC 1200 (Thermo) was used for proteomics measurements. For each fraction, ∼3 µg of peptides was dissolved in 5% ACN, and 5% FA before being loaded onto an in-house 100-μm capillary column packed with 35 cm of Accucore 150 resin (2.6 μm,150 Å). Peptides were separated and analyzed using a 180-min gradient consisting of 2%–23% ACN, 0.125% FA at 500 nL min^−1^ flow rate. A FAIMS Pro (Thermo) device was used for field asymmetric waveform ion mobility spectrometry (FAIMS) separation of precursors^62^, and the device was operated with default settings and multiple compensation voltages (−40V/−60V/−80V). Under each voltage, data-dependent acquisition mode was used for a mass range of m/z 400–1400 applying a top10 DDA method. Resolution for MS1 was set at 120,000. Singly-charged ions were not further sequenced, and multiply-charged ions were selected and subjected to fragmentation with standard automatic gain control (AGC) and 35% normalized collisional energy (NCE) for MS2, with a dynamic exclusion window of 30 s. Quantification of TMT reporter ions were performed using the multinotch SPS-MS3 method with 45% NCE for MS3, which is optimized for TMTpro-16 reagents.

##### Database searching

Raw files were first converted to mzXML and searched using the Comet algorithm on an in-house database search engine as reported previously.^63,64^ Database searching included all bacteriophage SPO1 (BPSP1) entries from UniProt, reverse sequences, and common contaminants. Peptides were searched using the following parameters: 25 ppm precursor mass tolerance, 1.0 Da product ion mass tolerance, and tryptic digestion with up to three missed cleavages. Methionine artificial oxidation (+15.9949 Da) was set as a variable modification. Carboxyamidomethylation (+57.0214637236) on cysteine was set as a static modification. For TMT-based experiments, TMTpro (+304.2071 Da) on lysine and peptide N-terminus were set as additional static modifications. The false discovery rate (FDR) was controlled as described previously described to <1% on peptide level for each MS run using parameters including XCorr, ΔCn, missed cleavages, peptide length, charge state, and precursor mass accuracy. Short peptides (<7 amino acids) were discarded, and the protein-level FDR was controlled to <1%.^64–66^

##### TMT reporter-based quantification

TMT reporter ions were used for quantification of peptide abundance. Each reporter ion was scanned using a 0.003 Da window, and the most intense m/z was used. Isotopic impurities were corrected according to the manufacturer’s specifications, and signal-to-noise ratio (S/N) was calculated. Peptides with summed S/N lower than 150 across 15 channels of each TMTpro16 plex (15 samples) or isolation specificity lower than 0.5 were discarded. Proteins were quantified by summing up the TMT S/Ns of peptides.

#### Cloning and plasmid construction

Codon-optimized sequences for SPO1 Acb4 and Acb4 homologs were synthesized as gBlocks (Integrated DNA Technologies) and assembled into the pETSUMO2 protein expression vector by Gibson assembly (NEB). Plasmids were transformed into the *E. coli* strain Top10, and construct generation was confirmed by whole-plasmid sequencing.

#### Recombinant protein expression and purification

SPO1 Acb4, Acb4 homologs, cGAS/DncV-like nucleotidyltransferases, cGAS-like receptors, and CBASS effector proteins were expressed and purified from *E. coli*. Synthetic gene fragments (Integrated DNA Technologies) were cloned by Gibson assembly (NEB) into a custom pET vector containing a 6×His-tagged N-terminal human SUMO2 fusion. Plasmids were transformed into *E. coli* BL21(DE3)-RIL cells (Agilent) and plated on MDG media (1.5% Bacto agar, 0.5% glucose, 2 mM MgSO_4_, 0.25% aspartic acid, and trace metals) with appropriate antibiotic selection (100 µg mL^−1^ ampicillin and 34 µg mL^−1^ chloramphenicol). Transformed colonies were used to inoculate a ∼30 mL starter culture and grown overnight for ∼17 hours in MDG media at 37°C with shaking (230 rpm). Expression cultures were inoculated with ∼15 mL of MDG starter culture in 1 L of M9ZB media (47.8 mM Na_2_HPO_4_, 22 mM KH_2_PO_4_, 18.7 mM NH_4_Cl, 85.6 mM NaCl, 1% casamino acids, 0.5% glycerol, 2 mM MgSO_4_, trace metals, 100 µg mL^−1^ ampicillin, and 34 µg mL^−1^ chloramphenicol) and grown to an OD_600_ of ∼2.5-3.0 prior to induction with 0.5 mM isopropyl-β-D-thiogalactoside (IPTG) for ∼15 hours at 16°C with shaking (230 rpm). After overnight expression, cultures were harvested by centrifugation at 3,500 × g for 20 minutes. The cell pellets were resuspended in 60 mL of lysis buffer (20 mM HEPES-KOH pH 7.5, 400 mM NaCl, 10% glycerol, 30 mM imidazole, and 1 mM DTT) and lysed by sonication to release soluble proteins. The lysate was clarified by centrifugation at 50,000 × g for 30 minutes and the resultant supernatant was purified by nickel affinity chromatography. Briefly, the lysate was poured over 8 mL of Ni-NTA Agarose (Qiagen) in a gravity flow column. The resin was washed sequentially with 20 mL of lysis buffer, 70 mL of wash buffer (20 mM HEPES-KOH pH 7.5, 1 M NaCl, 10% glycerol, 30 mM imidazole, and 1 mM DTT) and 35 mL of lysis buffer. The protein was eluted with 20 mL of elution buffer (20 mM HEPES-KOH pH 7.5, 400 mM NaCl, 10% glycerol, 300 mM imidazole, and 1 mM DTT). The eluate was treated with ∼200 µg of recombinant human SENP2 protease to facilitate cleavage of the 6×His-SUMO tag and dialyzed overnight at 4°C in 14 kDa molecular weight cutoff dialysis tubing (Ward’s Science). Proteins used for crystallography were dialyzed into dialysis buffer (20 mM HEPES-KOH pH 7.5, 250 mM KCl, and 1 mM TCEP), concentrated using a 10 kDa MWCO centrifugal filter (Millipore Sigma) then further purified by size-exclusion chromatography using a 16/600 Superdex 75 column (Cytiva). Proteins used for biochemical analyses were dialyzed into dialysis buffer, concentrated, aliquoted, flash frozen in liquid nitrogen and stored at −80°C until use in downstream assays.

#### Crystallization and structure determination

Crystals of the SPO1 Acb4–3′3′-cGAMP complex were grown using the hanging-drop vapor diffusion method for ∼9 days at 18°C. Recombinant Acb4 was diluted to 10 mg mL^−1^ in a buffer containing 100 mM KCl, 20 mM HEPES-KOH pH 7.5, and 1 mM TCEP. The diluted protein was incubated with 1 mM 3′3′-cGAMP (Biolog Life Science Institute) on ice for 10 minutes. The resultant protein mixture was allowed to equilibrate to 18°C and crystals were grown in 15-well trays (NeXtal) containing 400 μL reservoir solution and 2 μL drops. Drops were mixed 1:1 with purified protein and reservoir solution (18% PEG-4000, Tris-HCl pH 8.80, and 0.1 M MgCl_2_). Crystals were cryo-protected with reservoir solution supplemented with 24% ethylene glycol and 1 mM 3′3′-cGAMP (Biolog Life Science Institute) and harvested by flash-freezing in liquid nitrogen.

X-ray diffraction data were collected at the National Synchrotron Light Source II (NSLS2) beamline 17-ID-2 (FMX). Data were processed with autoPROC and Aimless.^67,68^ Experimental phase information was determined by molecular replacement using Phaser-MR in PHENIX and a model of the Acb4 structure generated using AlphaFold2.^69,70^ Model building was performed using Coot^71^ and refined using PHENIX. A summary of crystallographic statistics is provided in Table S2. All structural figures were generated using PyMOL (Version 2.5.4, Schrödinger, LLC).

#### Phylogenetic analysis of Acb4 homologs

Phage SPO1 Acb4 belongs to a previously uncharacterized protein family designated with the Pfam accession number PF13876.^44^ To construct the phylogenetic tree in Figure 5A, sequences belonging to the Acb4 Pfam were downloaded and aligned using MAFFT.^72^ The final aligned 1,331 sequences were used to construct a phylogenetic tree in Geneious Prime (v2024.0.3) using the FastTree plugin. iTOL (v6) was used to annotate the tree with taxonomic information extracted from NCBI.^73^ Full sequences and taxonomic information for the 1,331 analyzed Acb4 homologs are provided in Table S3.

#### *In vitro* reconstitution of CBASS effector function and inhibition

To determine the effect of Acb4 on CBASS effector activation, synthetic 3′3′-cGAMP was pre-treated with recombinant Acb4 and incubated with the CBASS nuclease effector protein *Burkholderia pseudomallei* Cap5.^14^ Briefly, 5 µM synthetic 3′3′-cGAMP (Biolog Life Science Institute) and 100 ng μL^−1^ pGEM9Z plasmid DNA was pre-incubated on ice for 30 minutes with 5 µM, 25 µM, or 50 µM recombinant Acb4. Pre-treatment reactions were prepared as 10× inputs and effector activation reactions were performed in 25 μL reactions containing 1 µM *Bp*Cap5 in a buffer consisting of 50 mM Tris-HCl pH 7.5, 25 mM NaCl, 5 mM MgCl_2_, and 1 mM TCEP. Effector activation reactions were incubated for 30 minutes at 37°C and reactions were terminated with the addition of 5 μL 6× DNA loading dye containing EDTA (NEB). To analyze Cap5 nuclease and DNA cleavage activity, 20 μL of the reaction was resolved on a 1% agarose gel run at 110V for ∼70 minutes in 1× TAE buffer and post-stained with ethidium bromide.

#### Phage challenge assays

Solid media phage challenge experiments were performed by spotting serial dilutions of high-titer phage stocks onto a lawn of bacteria harboring an active CBASS defense operon. Briefly, the CBASS operons *Yersinia aleksiciae* CdnE^19^ and *Escherichia coli* CdnG^14,21^ were cloned into an arabinose-inducible pBAD vector. Control plasmids were generated by inactivating the *Ec*CdnG CD-NTase^21^ (D82A/D84A) or deleting the transmembrane domain of the *Ya*CdnE effector. Cloned plasmids were transformed into *E. coli* BL21(DE3) cells (Agilent) and plated on LB agar with appropriate antibiotic selection. Liquid cultures were initiated from transformed colonies or glycerol stocks in ∼6 mL of LB media supplemented with appropriate antibiotic selection. Liquid cultures were grown for ∼5–6 hours at 37°C with shaking (230 rpm) and normalized to an OD_600_ of 0.40 in 16 mL of 0.5% top agar supplemented with 5 mM MgCl_2_, 5 mM CaCl_2_, 0.1 mM MnCl_2_, 0.02% (w/v) arabinose, and appropriate antibiotic selection. The top agar mixture was poured onto a 12 cm plate of LB agar containing 5 mM MgCl_2_, 5 mM CaCl_2_, 0.1 mM MnCl_2_, and appropriate antibiotic selection. Plates were allowed to solidify for ∼30 minutes at room temperature to allow for CBASS induction. High-titer phage stocks were serially diluted tenfold in SM buffer (50 mM Tris-HCl pH 7.5, 100 mM NaCl, and 8 mM MgSO_4_) and 3 μL droplets were spotted onto the bacterial top agar lawn. Plates were allowed to dry for ∼20–30 minutes at room temperature and incubated overnight at 30°C. Plates were imaged the following day using a BioRad ChemiDoc system.

#### Generation of phage T4 Δ*acb1* / Δ*acb2*::*acb4*

Recombinant T4 strains were generated using a Cas13a-based selection strategy, as described previously.^74^ We first designed gRNAs to target T4 *acb2* NC_000866.4 (nt 13-43 of NCBI gene ID: 1258619) and cloned them into the Cas13a-expressing vector pBA559, to generate the selective plasmid pBA559-*acb2*.^74^ To generate recombinant phages, we designed templates for homologous recombination consisting of either insertion mutations in T4 *acb2* that result in premature stop codons (Figure S3A) or SPO1 *acb4* flanked by ∼100 bp of homology upstream and downstream of the endogenous T4 *acb2* locus. These recombination templates were cloned into pGEM9z and used to transform *E. coli* Top10 cells (ThermoFisher). Colonies were picked into 2 mL of LB containing 100 μg mL^−1^ ampicillin and log scale cultures were infected with either WT T4 or T4 Δ*acb1* (published previously)^21^ and incubated at 37°C with shaking until culture clearance. Resulting phage lysates were passed through a 0.22 μM filter and stored at 4°C until used in selection. To select for recombinant T4 Δ*acb2*, T4 Δ*acb1/*Δ*acb2*, and T4 Δ*acb1/*Δ*acb2::acb4*, we transformed *E. coli* Top10 cells with pBA559-*acb2*, picked colonies, and grew overnight cultures in 2 mL LB with 34 μg mL^−1^ chloramphenicol. Top agar consisting of 0.6% agarose and 10 nM anhydrotetracycline (Cayman Chemical 10009542) was maintained at 42°C, and 500 μL overnight culture was added to 4 mL of top agar and plated on 20 mL of LB base agar containing 1.5% agarose and 34 μg mL^−1^ chloramphenicol. Following overnight incubation at 37°C, single plaques were picked and expanded at 37°C in liquid cultures of *E. coli* Top10 cells transformed with pBA559-*acb2* for a total of 3 rounds of plaque purification. PCR primers were designed to amplify the *acb2* locus and 1 μL of each phage stock was used as a template for a standard PCR reaction using Q5 polymerase according to the manufacturer’s instructions (NEB). PCR products were purified using the QIAquick gel extraction kit (Qiagen) and introduction of the desired mutations was confirmed by Sanger sequencing (Quintara Biosciences).

### QUANTIFICATION AND STATISTICAL ANALYSIS

Statistical analyses are outlined in the figure legends. Data are plotted with error bars representing the standard deviation (SD). For thermal proteome profiling experiments, all data analyses were performed in R (Version 4.2.1) unless stated otherwise. Pearson correlation analyses were either performed in GraphPad Prism (Version 10.2.2) or in R. Significance of cysteine exposure was determined with two-tailed Student’s t-tests for pairwise comparison.

