## Supplementary material for "A widespread family of viral sponge proteins reveals specific inhibition of nucleotide signals in anti-phage defense": Chang et al Table S2

**Table S2. Crystallographic Statistics, Related to Figures 3 and 4**

| | ***Bacillus* phage SPO1 Acb4–3′3′-cGAMP** |
|---|---|
| Resolution (Å)[a] | 83.73–2.06 (2.17–2.06) |
| Wavelength (Å) | 0.97934 |
| Space group | P 1 |
| Unit cell: a, b, c (Å) | 54.28, 56.50, 167.58 |
| Unit cell: $\alpha$, $\beta$, $\gamma$ (°) | 87.53, 89.02, 81.58 |
| Molecules per ASU | 20 |
| Total reflections | 430,233 (60,909) |
| Unique reflections | 119,318 (17,393) |
| Completeness (%)[a] | 97.8 (97.0) |
| Multiplicity[a] | 3.6 (3.5) |
| $I/\sigma I$[a] | 3.2 (0.6) |
| CC(1/2)[b] (%)[a] | 98.8 (38.9) |
| Rpim[c] (%)[a] | 11.7 (92.2) |
| Resolution (Å) | 83.73–2.06 |
| Free reflections | 1,411 |
| R-factor / R-free | 21.3 / 25.4 |
| Bond distance (RMS Å) | 0.002 |
| Bond angles (RMS °) | 0.497 |
| No. atoms: protein | 14,462 |
| No. atoms: ligand / ion | 900 |
| No. atoms: water | 887 |
| Average B-factor: protein | 32.62 |
| Average B-factor: ligand | 28.79 |
| Average B-factor: water | 30.71 |
| Ramachandran plot: favored | 99.27% |
| Ramachandran plot: allowed | 0.73% |
| Ramachandran plot: outliers | 0.00% |
| Rotamer outliers | 1.32% |
| MolProbity[d] score | 1.41 |
| Protein Data Bank ID | 9E4W |

[a] Highest resolution shell values in parentheses
[b] (Karplus and Diederichs, 2012)
[c] (Weiss, 2001)
[d] (Chen et al., 2010)
